# Rarely categorical and highly separable: how neural representations change along the cortical hierarchy

**DOI:** 10.1101/2024.11.15.623878

**Authors:** Lorenzo Posani, Shuqi Wang, Samuel P. Muscinelli, Liam Paninski, Stefano Fusi

## Abstract

A long-standing debate in neuroscience concerns whether individual neurons are organized into functionally distinct populations that encode information differently (“categorical” representations [1–3]) and the implications for neural computation. Here, we systematically analyzed how cortical neurons encode cognitive, sensory, and movement variables across 43 cortical regions during a complex task (14,000+ units from the International Brain Laboratory public Brainwide Map data set [4]) and studied how these properties change across the sensory-cognitive cortical hierarchy [5]. We found that the structure of the neural code was scale-dependent: on a whole-cortex scale, neural selectivity was categorical and organized across regions in a way that reflected their anatomical connectivity. However, within individual regions, categorical representations were rare and limited to primary sensory areas, and neuronal responses were instead very diverse. With theoretical arguments and empirical evidence, we demonstrate that the diversity of neural responses enables high-dimensional representations and, hence, high separability, allowing linear readouts to separate experimental conditions in many arbitrary ways. Indeed, when accounting for information that is actually encoded in each area, all cortical regions exhibit maximal separability. Our results indicate that cortical circuits prioritize diversity over categorical structure, supporting a computational regime geared toward high-dimensional, highly-separable neural representations.

---

Neural responses are often complex, depend on multiple variables (mixed selectivity [6]), and, in cognitive areas, they are very diverse and seemingly disorganized (see e.g., [3, 7, 8]). One of the challenges in neuroscience is finding meaningful “structure” in seemingly disorganized data. This structure has been traditionally investigated in single-neuron responses in the form of easily interpretable tuning curves (e.g., orientation selectivity in the visual cortex or place cells in the hippocampus). However, it is now possible to record from a large number of neurons, and new interesting structures are emerging, both in the statistics of the responses of individual neurons and at the level of population responses.

Here, we focused on both types of structure, how they are related to each other, and how they can be interpreted in terms of their computational implications. Consider the neural activity estimated in one particular time interval, recorded in different experimental conditions (e.g., in response to different sensory stimuli) [3, 9]. The responses of the different recorded neurons can be organized in a matrix (Fig. 1a), in which each row is the set of responses of a particular neuron to all the experimental conditions. This matrix defines two complementary spaces: the one spanned by the rows of the matrix, indexed by condition number, called the “conditions space” (shown in red in Fig. 1a), and that spanned by its columns, indexed by neuron number, called the “neural space” (shown in blue in Fig. 1a).

**Fig. 1.**
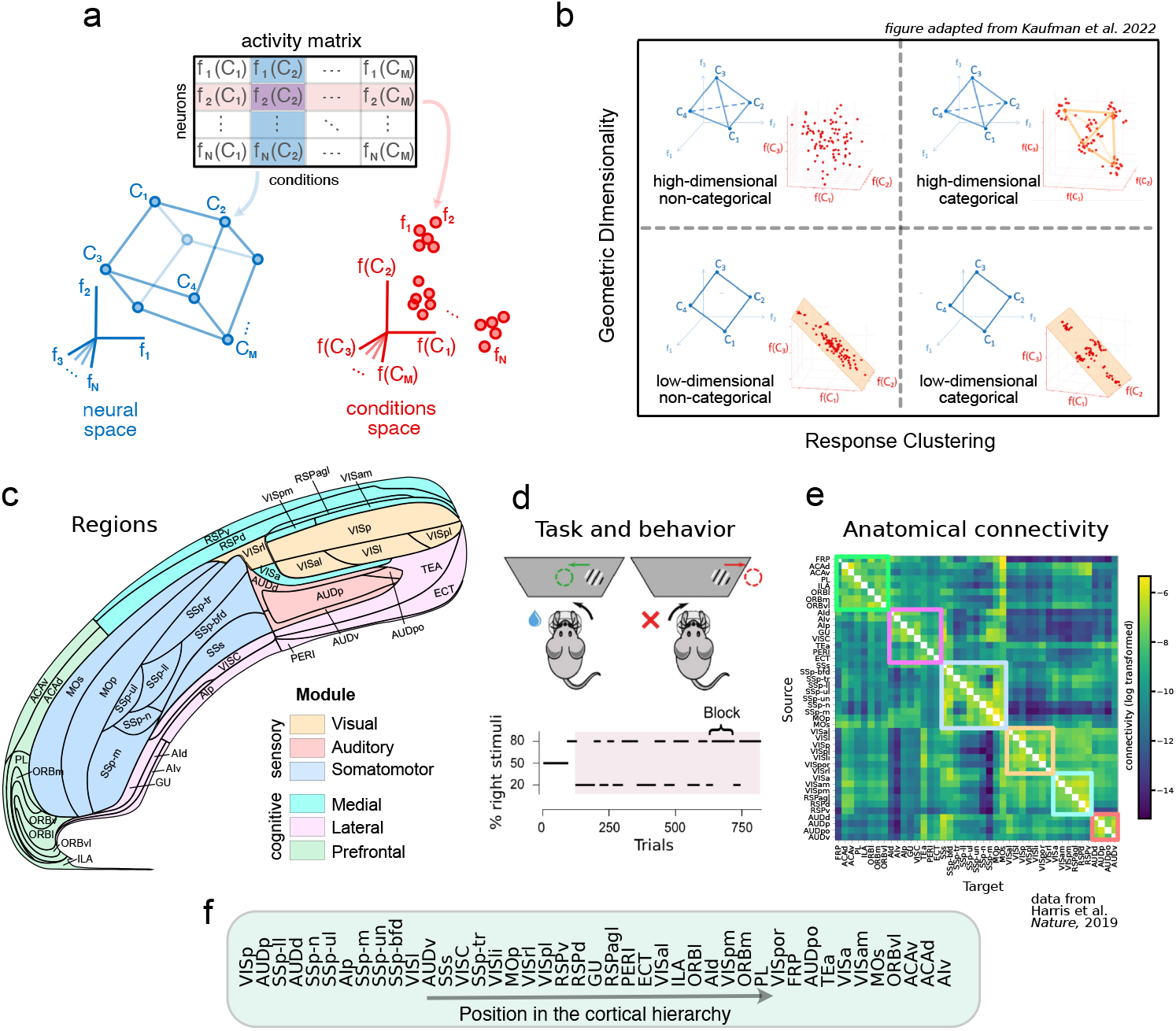
Conceptual framework and data structure: **(a)** The matrix of neural responses of *N* neurons in *M* experimental conditions can be analyzed in either its row space (conditions space, red) or column space (neural space, blue). The relative position of conditions in the neural space defines their “representational geometry”. If neurons are clustered in the conditions space, they define what is called a “categorical” representation [1–3]. **(b)** To what extent these two perspectives are related to each other is unclear, and neural populations could, in principle, occupy any portion of the clustering-dimensionality space. Image adapted from [3]. **(c)** Swanson flat map of the 43 cortical regions we analyzed. Data was recorded by the IBL consortium (IBL Brainwide Map dataset) using Neuropixel probes. After selecting neurons in the cortex, we were left with *∼* 14, 000 neurons from *∼* 180 recording sessions. **(d)** In the IBL task, mice need to rotate a wheel to move a visual stimulus toward the center of the screen. The stimulus appears left or right with an 80-20% biased probability in blocks of trials (bottom panel), adding “prior” contextual information to the task. **(e)** Region-to-region anatomical connectivity in the cortex, data from [5]. The colored boxes highlight six anatomical modules with dense intra-module connectivity. **(f)** Cortical hierarchy derived by the anatomical connectivity matrix of panel (e). The order was derived in [5] such that “source” regions, i.e., regions whose connectivity is unbalanced outwards, are mostly placed low, and “target” regions, i.e., regions whose connectivity is unbalanced inwards, are mostly placed high in the hierarchy. Additional details on how the hierarchy is computed can be found in [5].

In the conditions space, individual points represent the response profile of single neurons to the experimental conditions. If groups of neurons respond similarly in these conditions, they will form clusters (functional groups) in this space, defining what was called in [1] a *categorical* representation (Fig. 1a). For these representations, neurons with similar responses can be grouped together, forming a distinct category of neurons with specific functional properties. In non-categorical representations, neurons can have diverse responses whose distribution does not display clustering.

Categorical representations have been reported in the orbitofrontal cortex (OFC) of rodents [2, 10], and monkeys [11], while non-categorical representations were found in the rodent posterior parietal cortex (PPC) [1] and in reward-sensitive frontostriatal brain areas in monkeys performing a variety of economic decision-making tasks [12]. In several other articles [6, 13–16], the authors did not study explicitly whether the representations are categorical, but the observation of very diverse mixed selectivity neurons and high-dimensional representations suggests non-categorical representations (see also the Discussion). Whether, where, and how neural populations are subdivided into functional clusters is still an open debate [3].

In the neural space, spanned by the columns of the matrix, each point represents the population response to one experimental condition. The set of distances between all pairs of these points defines the *geometry* of the neural representations (Fig. 1a). Analyzing the representational geometry of population activity has been shown to provide insight into the brain’s encoding strategies and their computational implications for learning and flexible behavior [6, 14–17]. For example, a high-dimensional neural geometry confers flexibility to a simple linear readout [6–8], as the same representation can support a large number of different output functions. When the dimensionality is maximal, it is possible to retrain the readout to perform any task without changing the representation. High dimensional representations also maximize memory capacity [15], while low dimensionality is associated with a better ability to create abstract representations and generalize in novel situations [7, 14, 17]. How the dimensionality of population activity in the neural space is related to the clustering of response profiles in the conditions space is still an open question (Fig. 1b).

Here, we systematically analyzed the response profiles and the representational geometry in 43 cortical regions (Fig. 1c) in mice performing a decision-making task (IBL public Brainwide Map dataset [4], Fig. 1d), and related the functional properties of cortical areas with their anatomical properties as reported by the Allen Mouse Brain Connectivity Atlas ([5], Fig. 1e, f). As the neuronal responses have a complex temporal profile, we applied a reduced-rank regression model to characterize both the temporal and the selectivity components of the response profiles of individual neurons. We also employed different measures of embedding dimensionality [18] to characterize the geometry of neural representations and its computational implications.

We found that the selectivity profiles within individual areas are clustered (i.e., “categorical”) only in primary sensory areas. However, when multiple brain areas are considered, we observe clustering that reflects the brain’s large-scale organization, similar to what has been observed in the monkey and human brains (see e.g. [19]). The response profiles of neurons in different brain regions are different enough that a decoder can determine to which area a particular neuron belongs, with accuracy increasing for less-connected pairs of regions. Finally, we investigated the computational implications of the observed representations by studying the separability of the different experimental conditions, a measure of how many different classifications in the activity space can be solved by a linear readout [6]. We find that brain areas with more diverse neural responses (less structured) exhibit higher separability.

Together, these results reveal a systematic relationship between clustering and representational geometry, providing a new, large-scale perspective on neural selectivity and population coding across the cortex.

## Results

### Characterizing the response profiles of individual neurons

We first systematically studied the response profiles of individual neurons and compared their statistics across brain regions. To characterize the response profiles, we employed a linear encoding model to estimate the effect sizes of multiple behavioral covariates of interest on single-neuron responses. Specifically, we used a reduced-rank regression (RRR) model that predicts the activity of single neurons in response to changes in a set of variables that characterize each experimental condition and the behavior of the subject (Fig. 2a). The core component of the RRR encoding model is a small set of temporal bases, learned from data and shared across neurons to describe their time-varying responses (see Suppl. Fig. 1 for a schematic illustration of the RRR model and the Methods section for its mathematical formulation and for comparisons with previous regression models [4, 20, 21]). The sharing of the temporal bases significantly reduces the number of parameters and mitigates the issue of overfitting. Moreover, the resulting selectivity profile, composed of the estimated effect sizes of all variables, offers several advantages over the activity profiles defined by the mean responses to different experimental conditions, as it has better predictive power than the per-task condition trial average (Fig. 2b) and mitigates the issue of unbalanced or even missing conditions.

**Fig. 2.**
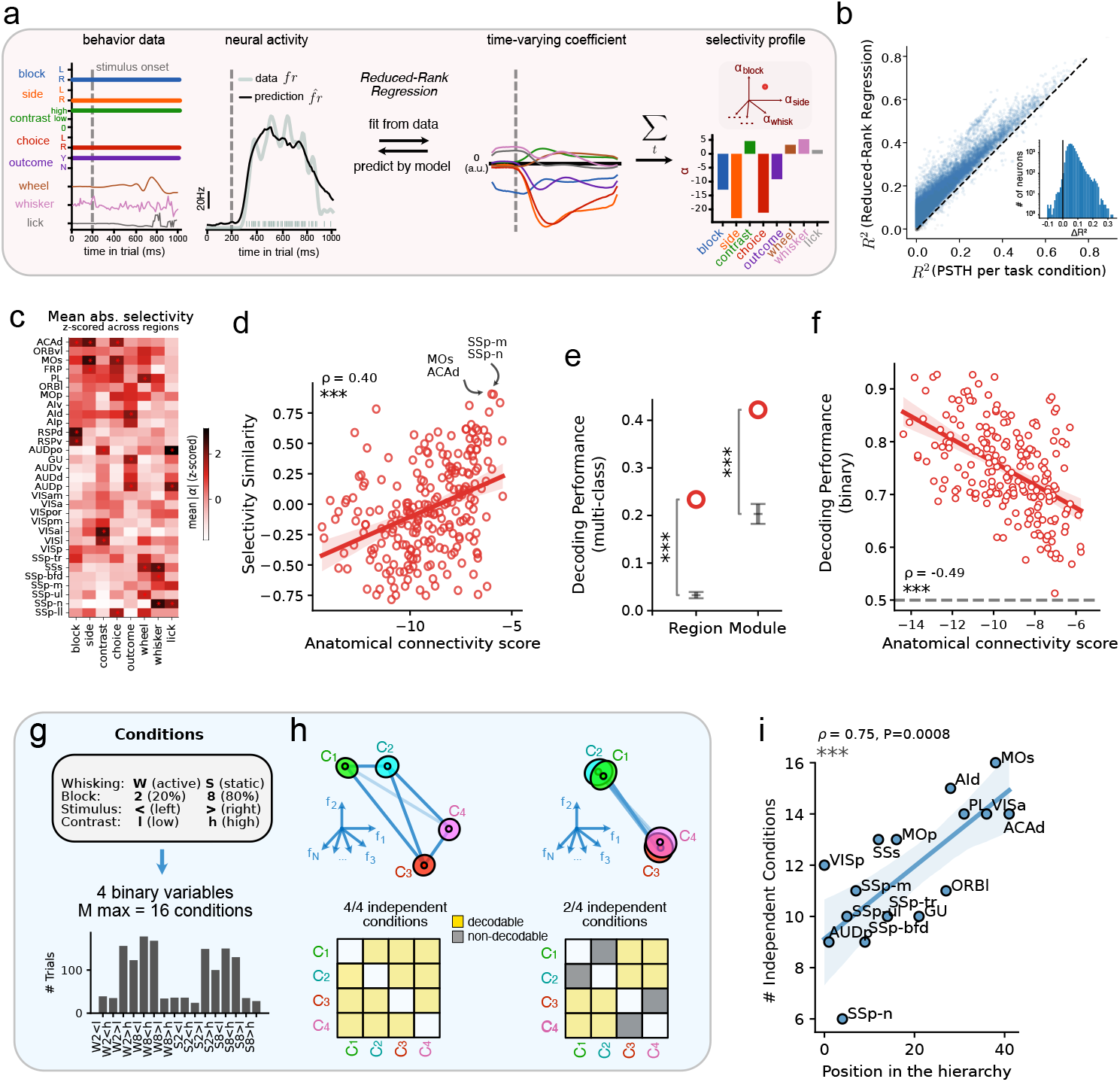
The reduced-rank regression encoding model as an effective and meaningful tool to estimate single neurons’ selectivity profiles. **(a)** Analysis pipeline for estimating single neurons’ selectivity profiles. A linear encoding model is first fitted to describe a single neuron’s temporal responses with respect to behavior data of interest (Methods). Selectivity (*α*) is computed as the sum of coefficients over time. Behavior data and neural activity of an example trial are shown for an example neuron, along with the estimated coefficients and selectivity. **(b)** Goodness-of-fit (three-fold cross-validated *R*^2^, see Methods) achieved by our encoding model versus computing the PSTH per task condition. The outperformance (Δ*R*^2^) is shown in the inset. **(c)** Average (absolute) selectivity profiles for neurons in the analyzed cortical areas. Areas are first grouped by their anatomical modules and then sorted by their hierarchy positions (see Fig. 1). The selectivity values are normalized per input variable (columns) for better visualization. The top three selective areas are marked with stars for each input variable. **(d)** Correlation between region-to-region functional similarity, measured as the cosine similarity between the average selectivity profiles in panel (e) and their anatomical connectivity (data in Fig. 1e). Each dot represents a pair of cortical areas. **(e)** Multi-class decoding performance of the region or module labels from the time-varying response profiles of individual neurons (panel a, third panel). Decoding performance is measured as the one-vs-one multi-class accuracy of a linear SVM, while the error bars indicate the distribution of decoding performance after random label shuffling. Module labels are defined as shown in Fig. 1, following the work of [5]. **(f)** The binary region-to-region decoding performance using the time-varying coefficients of single neurons inversely correlates with the anatomical connectivity (data in Fig. 1e) between the two regions involved. The dashed gray line indicates the chance level for binary decoding (50%). **(g)** Schematic for the analysis of independent conditions in the neural space. To select conditions, we chose the largest set of combinations of motor, sensory, and cognitive variables that are well represented in the behavior. The bottom panel shows an example of the number of trials for each condition in a session. **(h)** Schematic of two examples of geometries in the neural space with a different number of independent conditions (*M*_*IC*_ = 4 in the first case and *M*_*IC*_ = 2 in the second one). Independent conditions are found through an iterative algorithm based on the one-vs-one cross-validated decoding performance of all conditions; see Methods and Suppl. Fig. 5. **(i)** The number of independent conditions was found to vary across areas and increase along the hierarchy. \*\*\**p <* 0.001, \*\**p <* 0.01

We used this model to fit the response of each neuron (∼ 14, 000 neurons from 43 cortical regions) to 8 variables that describe the IBL decision-making task (Fig. 2a). In this task, subjects are shown a visual stimulus on one side of a screen and need to rotate a wheel to move the stimulus toward the center of the screen. They are given a water reward if they perform a correct wheel rotation. Stimuli are either shown on the left or right with a 50-50 balanced probability (“50-50 block”) or with an 80-20 imbalance (“left” or “right” block). In this analysis, we only used trials from unbalanced blocks to avoid confounds due to changes in neural signals over time, as the balanced block only appeared at the beginning of each session, and units may weaken or drop out over time.

The variables we used for fitting the model range from cognitive (left vs. right block) to sensory (stimulus side, contrast), motor (wheel velocity, whisking power, lick), and decision-related (choice, outcome). For each trial, we considered a time window of −0.2 to 0.8 seconds relative to the stimulus onset. After fitting the RRR model to predict trial-by-trial activity, each neuron is associated with a selectivity profile containing 8 ×5 = 40 coefficients (8 variables, 5 temporal bases). For each variable, we can then construct a time-varying coefficient by taking the weighted sum of the 5 temporal bases. These coefficients describe how sensitive the analyzed neuron activity is to each variable in trial time (Fig. 2a). We then summed these coefficients over time to estimate the total effect on neural responses and obtained a time-summed selectivity profile, which is an 8-dimensional selectivity vector (Fig. 2a, see also Suppl. Fig. 2 for examples of strongly selective neurons). Note that by normalizing the neuronal responses and input variables in the preprocessing steps (see Methods), we ensured that the unit-free coefficients 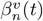 are not affected by the neuron’s mean firing rate or the inherently different scales in different input variables and can be compared directly across neurons, input variables, and time steps.

We first evaluated the predictive performance of our RRR model and compared it to the performance of the average firing rate per task condition (PSTH) computed in the same time interval used to fit the model, finding a significant improvement across the whole population of neurons (RRR: mean *R*^2^ = 0.16, PSTH: mean *R*^2^=0.09, *p* ≃ 0, Fig. 2b, see also Suppl. Fig. 3b for additional benchmark results and Suppl. Fig. 3c-f for visualization of example neurons). To prevent our results from being influenced by neurons that lack selectivity to any variable [4, 21], we included only neurons whose *R*^2^ exceeded a minimum Δ*R*^2^ threshold relative to a null model that does not incorporate variables as inputs (see Methods; 4617 neurons passed the threshold). As discussed in Suppl. Fig. 10, our results are robust to different values of this threshold.

**Fig. 3.**
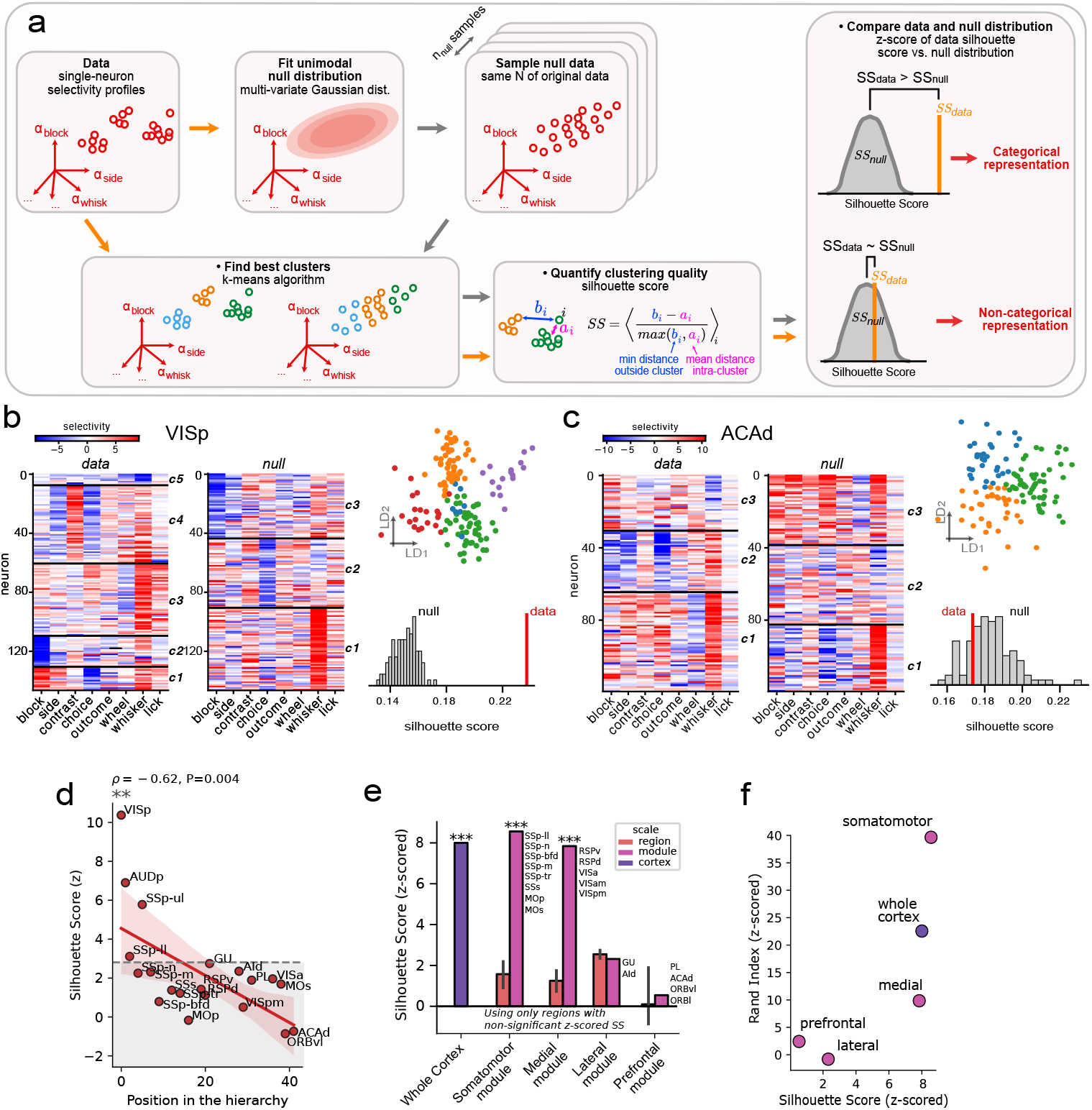
The clustering quality of areas varies across the cortical hierarchy and across anatomical scales. **(a)** Schematic plot of our pipeline to characterize categorical versus non-categorical representations. If representations are categorical, neurons will be clustered into multiple groups in terms of their response profiles, resulting in a silhouette score that is significantly larger than a non-categorical null model. The null model is a Gaussian distribution with a mean and covariance matrix matched to the data (see Methods and [1, 2]). For the non-categorical population, the silhouette score is comparable to the null distribution. **(b-c)** Examples of categorical (b) versus non-categorical (c) brain areas. On the left, the selectivity matrix of individual neurons is shown, with neurons sorted by the clustering labels and separated by black lines. The color indicates the selectivity value. In the middle, the selectivity matrix of an example null dataset is shown in the same way. The data silhouette score and its corresponding null distribution are shown at the bottom right, together with a reduced-dimensionality visualization (top right) obtained using linear discriminant analysis (individual points are neurons, colors refer to cluster labels). **(d)** Results of our categorical quantification pipeline in the selectivity profile space. For most cortical areas, the neuronal response profiles are very diverse, and the clustering structure is only present in primary sensory areas. The horizontal dashed line indicates the statistical threshold of significance (*p <* 0.05 with a Bonferroni correction for multiple comparisons). **(e)** Clustering quality by pooling neurons at different scales (purple: across the whole cortex, pink: across individual modules). Only neurons from non-clustered regions are considered. Red bars show the mean and standard deviation of the clustering quality of individual regions (that is, grouped data from panel d). **(f)** Cluster labels in well-clustered modules (medial, somatomotor) and across the whole cortex reflect the underlying anatomical organization. Similarity between anatomical labels and k-means clusters was quantified using the Rand index (RI), z-scored against a shuffled null model (see Methods), with higher z-scored RI values indicating a stronger correspondence between label sets. For anatomical labels, neurons pooled within a module were assigned area labels, whereas neurons pooled across the whole cortex were assigned module labels.

#### A hierarchical gradient of response timescales in the cortex

Previous and recent work has shown that neurons in regions higher up the cortical hierarchy exhibit progressively longer autocorrelation timescales, whether measured from spontaneous or task-related firing activity [22–25]. To determine if this trend extends beyond specific brain areas to the entire cortex, we analyzed the relationship between the autocorrelation timescales of neural responses to task variables – estimated using our RRR model (see Methods) – and the mouse cortical hierarchy defined by [5] from the Allen Institute Mouse Brain Connectivity Atlas. Our analysis complements prior work on intrinsic timescales of spontaneous activity using the IBL dataset [23]. We observed a significant positive correlation between a region’s hierarchical position and its average response timescale (Spearman correlation=0.62, p=0.002; Suppl. Fig. 4a,b). This finding aligns with earlier studies, reinforcing both the hierarchical organization of the cortex and the sensibility of our RRR model in capturing task-related activity.

**Fig. 4.**
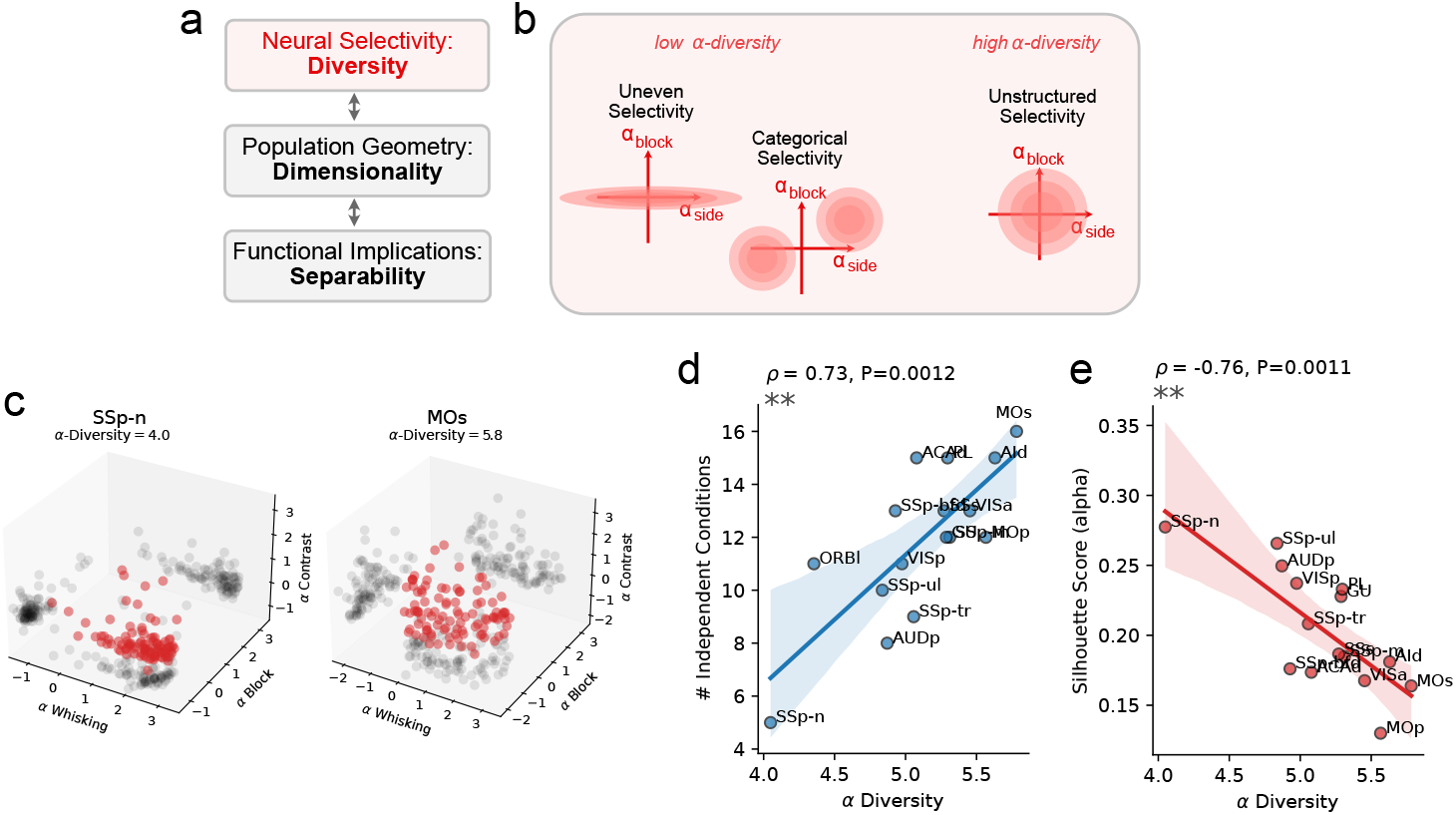
A unified measure of response profile diversity. **(a)** Conceptual framework, integrating previous findings and current results, illustrating how diversity in neural selectivity can enhance linear separability by increasing the dimensionality of the population geometry. **(b)** Schematic representation of the two main modes of structure observed in the data: uneven selectivity, where some variables or conditions are less encoded than others, and categorical selectivity, where neurons cluster into functionally distinct groups. **(c)** Examples of response-profile distributions for two cortical areas (SSp-n, MOs). For visualization, only three variables (whisking, block, contrast) are shown, while *α*-diversity is defined as the participation ratio of response profiles in the full 8-dimensional space of regression coefficients. SSp-n exhibits an elongated response profile dominated by whisking encoding (uneven selectivity), and has a low *α*-diversity (*≃*4.0). In contrast, MOs, a high-hierarchy region, displays a more uniform and diverse response distribution (*α*-diversity *≃*5.8). **(d)** Relationship between *α*-diversity and the number of independent conditions (*M*_*IC*_ ) across cortical regions. Higher *α*-diversity values were associated with a greater number of independent conditions (Spearman’s *ρ* = 0.73, *P* = 0.0012). **(e)** Relationship between *α*-diversity and the Silhouette Score in the *α*-diversity space. Regions with higher *α*-diversity showed lower clustering (Spearman’s *ρ* = *−*0.76, *P* = 0.0011).

### Not “everything everywhere”: response profiles reflect the anatomical organization of the cortex

Consistent with the dataset’s original paper [4], we found that many of the task-relevant variables are encoded in multiple brain regions. However, this does not mean that “everything is everywhere” in the sense that there are no differences in the response properties of different brain regions. Indeed, as we will show, we observed that the statistics of the response profiles of individual neurons are distinct to the point that we can guess the region a neuron belongs to with greater-than-chance accuracy.

The time-aggregated response profiles averaged across the neurons in each specific brain area are reported in Fig. 2c. To compare functional and anatomical properties of cortical regions, we computed the cosine similarity of the average selectivity profiles for all pairs of regions and compared these to the region-region anatomical connectivity reported in [5] (Fig. 1d). We found a significant positive correlation between selectivity similarity and anatomical connectivity (Spearman correlation = 0.40, *p* = 1.8 · 10^*−*10^, Fig. 2d), indicating that anatomically distant regions have more distinct average response profiles.

Encouraged by this result, we wondered whether selectivity profiles of individual neurons could be used to determine their region of origin. We thus labeled each neuron’s selectivity profile with its region of origin and used a cross-validated multi-class decoding approach to assess the extent to which this information could be decoded. We found that the region of origin could be decoded from individual neurons’ selectivity profiles with above-chance accuracy using both time-varying selectivity profiles (accuracy: 0.233; null: 0.033 ± 0.003; *p* ≃ 0; Fig. 2e) and time-summed selectivity profiles (accuracy: 0.123; null: 0.018 ± 0.002; *p* ≃ 0). Similarly, we successfully decoded the anatomical position of individual neurons on the larger scale of brain *modules* (accuracy: 0.422; null: 0.203 ± 0.010; *p* ≃ 0; Fig. 2e), as defined by [5], which proposed these region groupings based on anatomical connectivity and previously known functional properties. We then used the selectivity profiles of each neuron to compute the binary region-to-region decoding performance, finding that the decoding performance of a pair of regions inversely correlated with their anatomical connectivity (Spearman corr: -0.49; *p* = 3.7 · 10^*−*14^; Fig. 2f). Note that in all these cases, we decoded the brain region from the response profiles of individual neurons. The resulting decoding accuracy is perhaps surprisingly high, given the diversity of neuronal responses within each area, as discussed below.

We next wondered whether we could observe systematic differences across the cortical hierarchy in the coding properties of neural populations for task variables. We thus performed an analysis aimed at finding how many “independent” conditions a neural population encodes out of all possible combinations of task variables. We call this measure *M*_*IC*_. This measure is intimately related to the amount of information a neural population carries about the experimental task [26]. Importantly, it allows for non-linear combinations of task variables to emerge as encoded conditions, thus not requiring us to make strong assumptions about which variables to test.

We first defined a discrete set of conditions by using the combinations of 4 variables, which were chosen so that they span different categories (sensory, motor, and cognitive) and so that each condition was well represented in the behavior: whisking motion, block prior, stimulus contrast, and stimulus side (see Methods and Suppl. Fig. 10). Continuous variables (e.g., whisking motion) were binarized to make them consistent with the other binary task variables (e.g., block prior). By experimental design, variables like context and stimulus side are highly correlated in the biased blocks of trials that we analyzed. Hence, adding additional variables (such as choice or reward) was impractical due to the very limited number of trials for certain variable combinations (e.g., errors in context-side matched trials). This choice of variables defines a maximum number of independent conditions 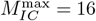 (Fig. 2g).

We then estimated *M*_*IC*_ for each region using an iterative algorithm based on the cross-validated decoding performance of a linear classifier trained to distinguish pairs of conditions in a 1-v-1 decoding test and grouping non-decodable conditions under a new, single label. The algorithm iterates this procedure until all labeled conditions are pairwise decodable from each other (Fig. 2h, see Methods and Suppl. Fig. 5). The number of independent conditions varied widely across different cortical regions, ranging from a minimum of *M*_*IC*_ = 6 (SSp-n) to a maximum of 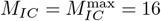 (MOs). *M*_*IC*_ increased significantly along the cortical hierarchy (Spearman corr: 0.77, *p* = 0.0005; Fig. 2i). Thus, cortical regions encode an increasing number of combinations of task variables as they move higher up the cortical hierarchy.

**Fig. 5.**
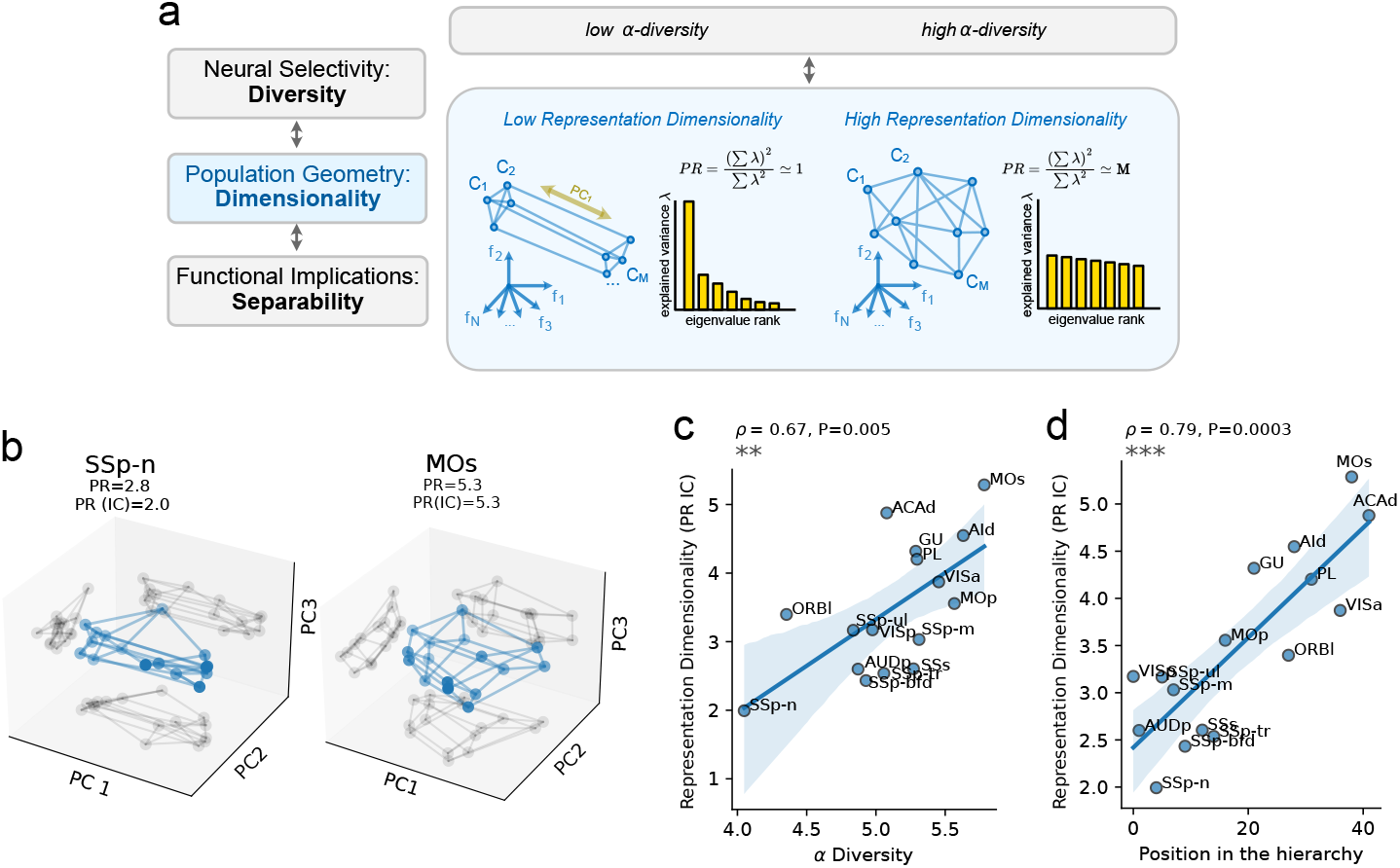
Relation between structure in neural response profiles and embedding dimensionality of neural representations. **(a)** Conceptual illustration of low-versus high-dimensional population geometries. When only a few eigenvalues dominate the covariance spectrum, the participation ratio (PR) is low, indicating that most variance lies in a small number of dimensions (left). When eigenvalues are more evenly distributed, PR is high, corresponding to a more isotropic geometry (right). **(b)** Example population geometries for two cortical regions (SSp-n, MOs) projected onto the first three principal components of the *M* = 16 centroids. The low-hierarchy, low-diversity area SSp-n exhibits reduced representation dimensionality compared with MOs. **(c)** Representation dimensionality is higher in areas with higher diversity of response profiles. **(d)** Representation dimensionality increases systematically along the cortical hierarchy.

Together, the double line of evidence based on single-cell response profiles and population decoding suggests that information is not encoded equally across cortical regions, but varies across the hierarchy in a way that reflects the large-scale anatomical organization of the cortex.

### Within regions, neural representations are rarely categorical

We then focused on the selectivity properties of single neurons within each area and investigated whether neurons cluster into specialized sub-populations (“categorical” selectivity) or whether information about task variables is heterogeneously distributed across the whole population (“non-categorical” selectivity).

As noted above, our RRR model associates a time-summed selectivity profile (8-dimensional selectivity vector) to each neuron. Thus, a population of neurons within a region can be visualized as a cloud of points in an 8-dimensional selectivity space (Fig. 3a). As a measure of clustering quality, we used the silhouette score (SS) [27] of clusters identified using k-means in this space. For a given neuron, the SS compares its mean distance to neurons within the same cluster with the distance to the nearest out-of-cluster points (Fig. 3a).

Fig. 3a describes our pipeline to compute the clustering quality of a neural population: first, we used k-means to find the clustering labels. The *k* parameter is chosen to maximize the SS. We then computed the SS for these clusters, which we call *SS*_data_. Importantly, across all the clustering analyses, we considered only clusters that are reproducible, i.e., that are not dominated by neurons from a single experimental session (see Methods). We then compared the SS observed in the data with a null model that is sampled from a single Gaussian distribution (which is unimodal, thus non-clustered by design) while preserving the mean, covariance structure, and the number of neurons of the original data. This null model is crucial for debiasing silhouette scores across varying dimensionalities (Suppl. Fig. 6) and was inspired by the one used for the ePAIRS test from [2]. By repeating many null model iterations, we could compare the *SS*_data_ value with a null model distribution of silhouette scores ({*SS*_null_}). As a measure of clustering quality, we took the deviation of the data point from the null distribution, measured with the z-score.

**Fig. 6.**
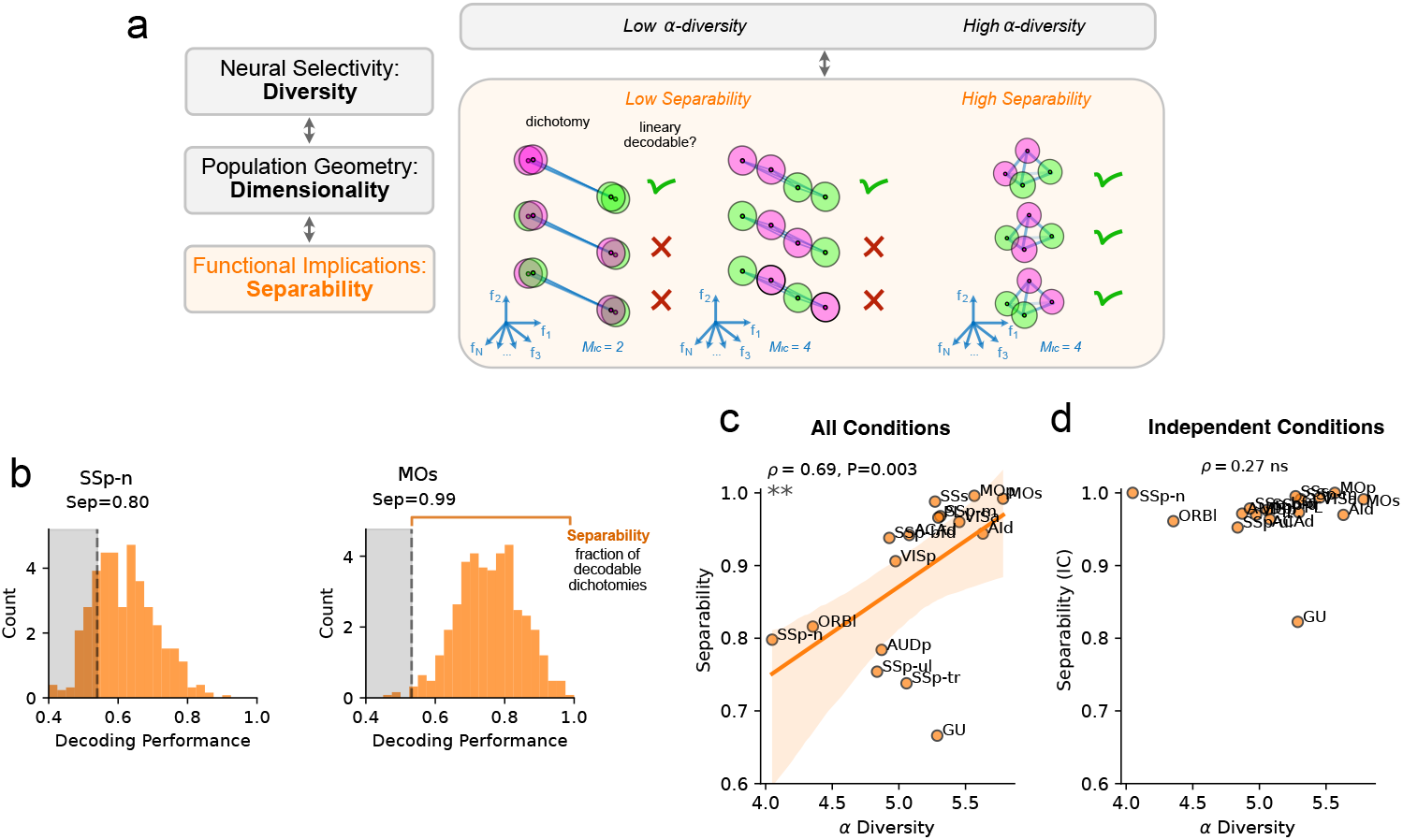
Diversity in neural response profiles is predictive of linear separability. **(a)** Illustration of three geometries of *M* = 4 conditions with different degrees of Separability, defined as the fraction of balanced dichotomies (represented by coloring schemes) that can be linearly separated in the activity space. Points represent conditions in the neural space, while clouds represent trial-to-trial noise. The geometry on the left has low Separability (1/3 decodable dichotomies) because of a lower number of independent conditions (*M*_*IC*_ = 2). The geometry in the middle has a maximal number of independent conditions (*M*_*IC*_ = 4), but still exhibits low Separability (1/3 separable dichotomies) due to the collinearity of conditions in the neural space. The geometry on the right has the maximum dimensionality (3-dimensional tetrahedron) and *M*_*IC*_, and supports maximal Separability (3/3 decodable dichotomies). **(b)** Distribution of cross-validated decoding performance of a linear classifier trained to decode *n* = 200 random balanced dichotomies of the *M* = 16 conditions defined in Fig. 2g, for SSp-n (left) and MOs (right). The dashed lines and shaded areas indicate the 99 percentiles of a shuffled null model for the decoding performance (see Methods). Separability is defined as the fraction of dichotomies for which the linear decoding performance exceeds this threshold; **(c)** Separability, computed across cortical regions, using the entire set of *P* = 16 conditions from Fig. 2g, is higher for regions with diverse neural responses **(d)** Conversely, Separability of independent conditions (Fig. 2i) is always maximal (with one exception in the GU area) across all cortical regions.

We analyzed the clustering structure of selectivity profiles in all the recorded cortical regions. To avoid small-size artifacts, we included only regions that have at least 50 neurons after our *R*^2^ threshold was applied. In Fig. 3b, c, we show two examples of clustering results, one in a region that was found to be clustered (VISp) and one in a region that was found to be non-clustered (ACAd). To visualize the putative clusters found by k-means, we grouped neurons according to their cluster labels and sorted them according to their individual SS scores (highest at the top). As shown in Fig. 3b, the k-means algorithm clustered VISp neurons based on the block prior (c1, c2), whisking power (c3, c5), and a combination of side, contrast, and choice (c4). See also Suppl. Fig. 7 for the clustering results of all cortical regions.

The visual structure in these representations can be misleading, because apparent patterns may be driven by only a small number of neurons with strong, similar selectivity. Such local features do not necessarily reflect the overall population distribution. For example, a handful of strongly selective neurons is fully consistent with a globally unstructured Gaussian distribution. Indeed, we observe similar visual structure in a null-model sample after sorting (see c1 and c3 in Fig. 3b, middle panel). This example showcases the necessity of comparing the silhouette score against a null model distribution to obtain a meaningful measure of clustering quality. In the case of VISp, the primary visual area, the data clusters were found to have a higher SS than the null model (Fig. 3b, right panel), indicating that neural selectivity is more categorical than a randomly distributed Gaussian null model. This was not the case for ACAd, a prefrontal area (Fig. 3c), despite some apparent clusters in the selectivity profiles, mainly driven by a combination of stimulus side and choice (c2) and whisking (c1, c3).

When applying this pipeline to all cortical areas, we found that only a few areas passed the statistical threshold of significance (*p* < 0.05 with a Bonferroni correction for multiple comparisons). The primary visual (VISp) and auditory (AUDp) areas, as well as the upper limb region of the primary somatosensory cortex (SSp-ul) showed a highly significant silhouette score, while the lower limb region (SSp-ll) and the gustatory cortex (GU) sit close to the threshold. To test which variables contributed to the quality of clusters, we removed each variable individually and measured the drop in silhouette score caused by this removal. As shown in Suppl. Fig. 8, the most important variables for determining the clustering quality varied across areas and were typically multi-modal, spanning cognitive, movement, and sensory variables (VISp: block, whisking, contrast, and choice; AUDp: reward and whisking; SSp-ul: wheel and choice).

We also observed that clustering quality, measured by the z-score of the silhouette score against the null model, was significantly larger than that of the unstructured null model only in early sensory areas (Spearman correlation with position on the hierarchy: -0.62, p=0.004; Fig. 3d), suggesting that the neural code becomes more randomly mixed along the sensory-cognitive hierarchy. This negative trend was consistently found across several analysis choices such as varying the *R*^2^ threshold for neuron inclusion (Suppl. Fig. 10b), changing the clustering algorithm (Leiden algorithm [28], Suppl. Fig. 10c), and considering the specific temporal profiles of the time-varying selectivity coefficients on top of the sum across trial time (Suppl. Fig. 10d).

One potential weakness of the RRR model-based analysis approach is that it makes explicit assumptions about what variables are represented in the neural activity [2]. Therefore, we repeated the clustering analysis using the activity profiles in the conditions space, i.e., the space of trial-average firing rate of neurons in response to experimental conditions. For this analysis, we used the 16 conditions identified above (Fig. 2g) by combining four binarized task variables spanning sensory, motor, and cognitive modalities (see Methods and Suppl. Fig. 10). Next, we quantified the degree of clustering in the conditions space, with a pipeline inspired by [2]: we pre-processed the mean firing rates (z-score and dimensionality reduction), quantified the structural properties of neurons in the resulting pre-processed conditions space, and compared them to an uni-modal, non-clustered null model fitted from the data (see Methods). However, our pipeline differs from [2] in that the null distribution is fitted on the original data instead of the preprocessed firing rates (Suppl. Fig. 9a). This choice avoids the strong assumption of normality of the preprocessed data of the original method, showing a better fit to the data (Suppl. Fig. 9b-d). Once again, we observed a negative correlation between the hierarchy and the clustering quality in this 16-dimensional conditions space (Spearman correlation: -0.52, *p* = 0.039; Suppl. Fig. 10f), with only a few regions (VISp, AUDp, SSp-n, MOp) showing more clustering than the null model.

Finally, we used ePAIRS [2] to check whether there is any additional structure in the selectivity profiles that our pipeline might not capture: ePAIRS tests for deviations from a randomly mixed selectivity distribution without explicitly conducting clustering analysis. ePAIRS also yielded few brain regions passing the test for being categorical (VISp and MOs, Suppl. Fig. 10g), but showed no correlation with the hierarchy (Spearman correlation: -0.19, *p* > 0.05).

Together, these results provide systematic evidence that few cortical areas represent the experimental variables with a categorical representation, and that these structured areas are typically positioned at the lower end of the hierarchy.

### Mesoscale clustering in somatomotor and medial modules

The analysis of response profiles and independent conditions of Fig. 2 shows that, at a larger scale, the brain is functionally organized. When considering multiple brain regions together, we therefore expect to observe more categorical representations, as each region contains one or more neuronal populations with specialized response properties.

We applied our clustering pipeline to neural populations pooled across cortical modules defined in [5]: visual, auditory, somatomotor, medial, lateral, and prefrontal (Fig. 1c,e). To specifically test whether modules that were not categorical at the single-area level would show clustering on a larger scale, we excluded regions that had already greatly exceeded the clustering significance threshold (VISp, AUDp, and SSp-ul). Because the only analyzed regions in the visual and auditory modules (VISp and AUDp) were already clustered, our mesoscale analysis focused on four modules: somatomotor, medial, lateral, and prefrontal. To prevent overrepresentation by large regions, we selected the 100 best-encoded neurons from each area before pooling. This analysis revealed categorical representations in the somatomotor and medial modules (somatomotor: *z* ≃ 8; medial: *z* ≃ 8; Fig. 3e), whereas the prefrontal and lateral modules were not categorical (Fig. 3e). A more detailed analysis of the variables driving the module-level clustering is provided in Supplementary Fig. 11. To further assess how functional clusters relate to anatomical segregation, we computed the Rand Index (RI), which quantifies the similarity between clustering results and area labels (Fig. 3f), for all cortical modules. For the clustered modules (somatomotor and medial), the RI was significantly above chance (Fig. 3f, Suppl. Fig. 11), highlighting a structural correspondence between clusters found by our algorithm in the selectivity space and anatomically-defined regions.

Finally, when analyzing a single population spanning the entire cortex, representations were found to be categorical (*z* ≃ 8, Fig. 3e). In this case, nearly all variables contributed to cluster formation (Suppl. Fig.11b), and cluster labels significantly overlapped with module labels (Suppl. Fig. 11a,b).

This analysis provides evidence that the brain is globally organized and that functional specialization relates to the anatomical organization of the cortex.

### The functional implications of diversity in neural selectivity profiles

The results of Fig. 2i and Fig. 3d indicate that neural selectivity profiles are highly diverse in all brain areas analyzed and become increasingly diverse along the cortical hierarchy. What could this diversity be useful for?

In the following sections, we argue that high diversity is important for achieving high-dimensional representations in the neural space, which, in turn, enables better separability (Fig. 4a). We first define a measure of diversity in neural response profiles that captures the structure observed in the data (“*α*−diversity”). Using empirical and theoretical arguments, we then link single-neuron response diversity to the embedding dimensionality of population representations in the neural space (“Representation dimensionality”). Finally, we show that neural response diversity is directly related to the number of dichotomies of conditions that can be decoded by a cross-validated linear classifier (“Separability”), thereby connecting the structural properties of single-neuron selectivity to the functional properties of population-level representations.

### Characterizing structure in the selectivity space

The results of Fig. 2 and Fig. 3 suggest that neural selectivity is structured in two ways: first, it is *uneven* (“not everything everywhere”) as certain task variables are more strongly represented than others (Fig. 4b). This is revealed both by the diversity of the average linear regression coefficients across areas (Fig. 2c) and by the fact that the number of encoded conditions increases along the hierarchy (Fig. 2i). Second, it can be *categorical*, which is captured by the silhouette score (Fig. 3d). The silhouette score quantifies how well neurons cluster in functionally distinct subpopulations (Fig. 4b). Both forms of structured selectivity are more prevalent in sensory areas, and representations become more diverse up the cortical hierarchy.

To account for both forms of structure, we introduce a new measure of diversity, which we call *α*-diversity. This is defined as the participation ratio in the space of linear regression coefficients (*α*s, Fig. 2a), determined above using reduced rank regression. The participation ratio quantifies the number of dimensions spanned by a distribution of points in a high-dimensional space (see e.g. [26, 29]). Diverse responses are maximally unstructured when their distribution is an isotropic Gaussian. This distribution spans all dimensions of the *α* space uniformly, resulting in a high participation ratio. Any structure in the selectivity profile would make the responses more correlated, decrease the participation ratio, and hence decrease the *α*-diversity. In Fig. 4c, we show the distribution of response profiles for two areas (SSp-n, MOs). For visualization purposes, only the responses to three variables (whisking, block, contrast) are shown, but note that *α*-diversity is computed in the 8-dimensional space of all regression coefficients. SSp-n, a low-hierarchy area, has an elongated distribution across one dominant variable, whisking, resulting in a highly uneven selectivity distribution and a low *α*-diversity (≃4.0). On the contrary, MOs, a high-hierarchy area, shows a highly-diverse unstructured distribution (*α*-diversity ≃5.8).

Using synthetic explorations, we show that this measure directly captures variations in both categorical and uneven selectivity (Suppl. Fig. 12). This relation was indeed verified in the data, as *α*-diversity was directly correlated to the number of independent conditions (Spearman corr: 0.73, P=0.0012; Fig. 4d) and negatively correlated with the Silhouette Score in the *α* space (Spearman corr: −0.76, P=0.0011; Fig. 4e - although note that uneven selectivity also contributes to a higher silhouette score - see Suppl. Fig. 6).

Thus, *α*-diversity provides a unified quantitative measure for characterizing neural selectivity, encompassing categorical and uneven forms of structure, as well as potentially more complex properties that fall outside these simplified models.

### Diversity in neural selectivity enables high-dimensional representational geometries

As argued above, high-dimensional representations have been shown to be important for separability, flexibility, and memory capacity. Here, we investigate how structure in single neuron response profiles directly affects the dimensionality of representational geometries.

We consider the *Representation dimensionality* of a set of patterns of neural activity, which again can be expressed as the participation ratio (notice that the PR that we considered in the previous section refers to a different space, the space of the *α* coefficients). The participation ratio will be low if a few eigenvalues dominate (suggesting that only a few dimensions explain most of the variance, see Fig. 5a, top-left panel) and high if the eigenvalues are more evenly distributed (Fig. 5a, top-right panel).

To quantify the representation dimensionality, we computed the participation ratio of condition centroids in the *N*-dimensional neural space. We evaluated this measure using both the full set of *M* = 16 conditions defined in Fig. 2g (PR) and the subset of *M*_*IC*_ independent conditions (PR_IC_), to account for differences in noise level and condition discriminability across conditions. The latter was taken as the definition of representation dimensionality (this choice does not qualitatively impact our results, see Suppl. Fig. 15).

In Fig. 5b, we illustrate the representational geometry of the same two regions analyzed above, SSp-n and MOs, projected onto the first three principal components of the centroid distribution. As shown, SSp-n, a low-diversity, low-hierarchy area, exhibits a reduced representation dimensionality (PR_IC_≃2.0) compared to MOs (PR_IC_≃5.3), suggesting a link between higher diversity and higher representational dimensionality that was consistently observed across cortical regions (*α*-diversity vs. representation dimensionality; Spearman’s *ρ* =0.67, P= 0.005; Fig. 5c).

To understand this relationship quantitatively, we temporarily shift our focus from the space of regression coefficients (*α* space) to the *condition* space, defined by the average responses of single neurons to individual experimental conditions rather than task variables. Although these two spaces are linked in a non-trivial manner, structural organization in one is typically reflected in the other (for instance, VISp and AUDp are clustered both in the alpha space and in the conditions space - see Fig. 3d and Suppl. Fig. 10f).

The crucial intuition behind the relationship between structured selectivity and representation dimensionality is that conditions in the neural space (representational geometry) and neurons in the conditions space have *the same dimensionality* as they are, respectively, the column- and row-space of the same activity matrix (see Fig. 1a and Methods). Thus, when single-neuron responses are structured, their response profiles will be correlated, resulting in a lower-dimensional space in the conditions space and, in turn, reducing the representation dimensionality of population representations.

How does this intuition translate into a quantitative relationship in the two modes of structure we observed in the data? The relationship between *uneven* selectivity to conditions and representation dimensionality is straightforward: when certain conditions are more strongly encoded than others, the cloud of neural responses in condition space becomes elongated along specific axes, and the participation ratio decreases. Thus, in the absence of additional structure, a larger number of independent conditions will increase the representation dimensionality - a relation that was indeed observed in the data (M_IC_ vs. PR_IC_: Spearman’s *ρ* = 0.72, P= 0.0015; Suppl. Fig. 15).

In contrast, the quantitative relationship between *categorical clustering* and representation dimensionality is less straightforward. We can gain a qualitative intuition by considering a scenario in which the response profiles of neurons are organized into *k* < *M* small, well-defined clusters whose centers are randomly positioned in the *M*-dimensional conditions space (Suppl. Fig. 13a, bottom left). In the limit of small clusters, we have that all the neurons in each cluster are equivalent to a single neuron, and hence the effective number of neurons (i.e. the number of points in the conditions space) is equal to the number of clusters. When the number of conditions is large enough, the dimensionality is bounded by the number of points in the space, and hence it cannot be larger than the number of clusters. If we were to perform a PCA analysis on these response profiles, we would find that a high portion of the variance can be explained by the first *k* principal components, one for each cluster, yielding a low participation ratio *PR* ∼ *k* < *M*. On the contrary, non-categorical representations allow (but do not constrain) geometries to be high-dimensional (Suppl. Fig. 13a, right). To quantitatively study more intermediate scenarios, we developed a mathematical model that allowed us to analytically derive the Representation dimensionality in the neural space as a function of properties of categorical clusters: their number *k*, their diversity *δ*, and the number of conditions *M* (see Methods and Suppl. Fig. 13b). The derived formula shows that greater categorical representations (smaller *δ*) reduce the dimensionality of neural representations, predicting an inverse relation between Silhouette Score and Representation dimensionality (Suppl. Fig. 13b). This prediction was indeed verified on the data, both qualitatively (Representation dimensionality vs. SS in the conditions space: Spearman Corr=−0.77, *P* ≃ 0.0004, Suppl. Fig. 13c) and quantitatively (predicting Representation dimensionality from clustering properties and vice-versa, see Suppl. Fig. 13d,e).

As both categorical and uneven structure are more present in sensory than in cognitive areas (Fig. 2i, Fig. 3d), we expect that the Representation dimensionality will *increase* across the hierarchy. This was indeed observed in the data (Representation dimensionality vs. position in the hierarchy: Spearman’s Corr=0.79, *P* ≃ 0.0003, Fig. 5d). This result is similar to what was observed in the visual hierarchy in different experiments [30].

### Diversity in neural response profiles is predictive of linear separability

So far, we have shown that diversity in selectivity profiles and dimensionality of representational geometry are related to each other. How do these quantities translate into *functional* properties of neural representations? Several studies have shown that representations characterized by a high *embedding dimensionality* [9] in the neural space are important for cognitive flexibility [6, 7, 31] or high memory capacity [15]. All these studies are based on Cover’s theorem [32], which states that points in general position in a high-dimensional space are linearly separable in all possible ways. In other words, a linear classifier can partition the points (corresponding here to different experimental conditions) into any two arbitrary groups (“dichotomy”). Although originally formulated for noiseless settings, the theorem’s core intuition extends to noisy data. Indeed, Support Vector Machines with non-linear kernels, which are successfully applied to real-world noisy data, exploit the same principle: non-linear transformations increase the dimensionality of the feature space, thereby enhancing separability [33].

Here, we define Separability as the fraction of balanced dichotomies (different ways of dividing the conditions into two equal-sized groups) that are linearly separable. A dichotomy is linearly separable if the cross-validated performance of a linear decoder is larger than that of a null model in which the labels are shuffled. As we consider the cross-validated performance, trial-by-trial variations are unlikely to make neural representations separable. At the same time, noise can strongly affect separability (e.g. strong noise along the coding direction can highly decrease decoding performance).

In Fig. 6a, we show three schematic examples of how this notion of separability applies to different geometries. Each dot corresponds to a condition in the neural space, and the shaded area represents trial-to-trial variability (noise). In the first geometry (Fig. 6a, left panel), only 2 out of 4 conditions are independent. This low *M*_*IC*_ implies a low dimensionality and has a direct impact on Separability: only 1/3 of the dichotomies are linearly separable. However, Separability can also be low when the number of independent conditions is maximal. This is the case of the “collinear” geometry of Fig. 6a, central panel, where the centroids are positioned along a single axis. This low-dimensional geometry makes it impossible for a linear classifier to separate all dichotomies that are not defined along the linear axis, such as conditions in the middle versus conditions on the borders, resulting in a low Separability (= 1/3). Finally, the high-dimensional, unstructured tetrahedron in Fig. 6a, right panel, allows for all dichotomies to be linearly decoded, and Separability is maximal (= 1). A systematic analysis using synthetic data extends this intuition to larger dimensionalities, showing that Separability increases with the dimensionality of the geometry and saturates when the dimension is high enough (∼ *M*/2, see Suppl. Fig. 14c,d). As a complementary measure, we also studied Average Decodability (AD), introduced in [14] with the name “shattering dimensionality” and defined as the mean cross-validated decoding performance across all dichotomies (Suppl. Fig. 16). AD similarly increases with the representation dimensionality, however unlike separability, it does not easily saturate (Suppl. Fig. 14d).

As argued above, diversity is important for dimensionality. We thus expected *α*-diversity, Separability, and AD to be directly related to one another. We first computed Separability and AD in the synthetic model of uneven and categorical selectivity, showing that, for both types of selectivity structure, high neural diversity is related to high Separability (Suppl. Fig. 14b). We then studied this relation in real data. In Fig. 6b, we show the distribution of cross-validated linear decoding performance across dichotomies for the same two regions of Fig. 4 and 5, SSp-n and MOs. For each region, we trained linear SVMs on 200 randomly sampled dichotomies of conditions and measured their cross-validated decoding performance. The resulting distributions show that MOs achieves systematically higher decoding accuracy than SSp-n, resulting in higher Separability (MOs: Sep= 0.99; SSp-n: Sep= 0.80) and AD (MOs: AD= 0.76; SSp-n AD= 0.62), consistent with the prediction that higher neural diversity, and thus higher dimensionality, supports increased linear separability.

We then tested this relation across all cortical regions. As predicted, both Separability and Average Decodability (AD) increased systematically with *α*-diversity (*α*-diversity vs. Separability: Spearman’s *ρ* = 0.69, *P* = 0.003, Fig. 6c; *α*-diversity vs. AD: *ρ* = 0.66, *P* = 0.006, Fig. 16b), in accordance with our hypothesis that regions with more diverse single-neuron response profiles exhibit more linearly separable population codes. The analysis was performed using all *M* = 16 conditions defined in Fig. 2g, thereby capturing both categorical and uneven forms of selectivity.

We next asked how this relation changes when dichotomies are defined over *independent* conditions only, that is, when the effects of uneven selectivity are minimized. As shown in Suppl. Fig. 14d using synthetic data, separability is severely limited in low-dimensional geometries, even when all pairs of conditions are mutually separable (maximum independent conditions). Thus, by focusing on independent conditions, we aimed to isolate the contribution of geometric dimensionality to separability.

When analyzing Separability and AD for independent conditions in the data, we found that these quantities no longer correlated with *α*-diversity (Fig. 6d, Suppl. Fig. 16c). Separability was found to be near maximal (≥ 0.95) across nearly all cortical areas, with the sole exception of the gustatory cortex (Fig. 6d, Suppl. Fig. 17). Thus, once overlapping conditions are accounted for, cortical representations are not constrained by a low-dimensional geometry (as in Fig. 6a, middle panel) but instead occupy high-dimensional, unstructured spaces that guarantee high separability (as in Fig. 6a, right panel).

These results show that, although cortical regions differ in the number of encoded conditions and the degree of clustering, they represent those conditions with a dimensionality sufficiently high to achieve maximal linear separability.

## Discussion

The brain has a clear anatomical organization, and, not too surprisingly, we observe that it is reflected in the organization of the response profiles of individual neurons in different brain areas (functional organization). However, when one looks at the neurons within each brain area, only the responses of some primary sensory areas seem to be organized into functional clusters (categorical representations). The diversity of responses observed across brain areas plays an important computational role as it is related to the embedding dimensionality of the representations, which, in turn, determines their separability. We showed in a simple model how these different aspects of the representations are related to each other: structured selectivity limits the dimensionality of the representational geometry in the neural space, making diversity important for high-dimensional representations. High dimensionality enables simple linear readouts to perform a large number of input-output functions, or in other words, to separate the conditions in many different ways.

Finally, all our analyses revealed that several aspects of the representational geometry and the statistics of the neuronal response profiles vary in a systematic way along the cortical hierarchy: in particular, clustering decreases with position in the hierarchy, and the Representation dimensionality increases, along with the number of independent conditions. All these results show that the structure in the response profile space is related to the geometrical structure of the activity space. This relation can help us understand the computational implications of the neuronal response properties. All the analyses we performed can be applied to non-cortical regions, which is one of our future directions.

### Categorical representations

We observed that in most individual brain areas, the representations are non-categorical, whereas when the entire brain is considered, the representations are categorical. It is important to briefly discuss the meaning of this result: according to our definition, a representation is categorical if it has a silhouette score that is significantly different from that of the null model. The null model is a multivariate Gaussian distribution of neural response profiles. This means that any deviation from the null model distribution will be considered non-categorical, including distributions in which there are no identifiable categories of neurons (e.g., the continuous but structured representations observed in [34]). Our definition of non-categorical includes only a relatively narrow class of unstructured distributions, which makes our results even more surprising. Essentially, the primary structure observed in most individual brain areas is that the multivariate Gaussian distribution can be elongated along specific directions, which differ for each brain area. Notice that in the past, some investigators (notably the Freedman group [35, 36]) investigated categorical representations. However, in their case, ‘categorical’ refers to categories of stimuli, not to categories of response profiles, as we defined them here. It is possible that their representations are categorical in our sense, but to assess this, we would need to perform additional analysis. They show that some neurons are highly selective for categories of stimuli. The existence of these specialized neurons does not necessarily mean that the representations are categorical: these specialized neurons could simply be the neurons at the tail of a Gaussian distribution.

### Single neuron responses and representational geometry

Single neuron response properties and the geometrical structure of the activity space are related to each other. In particular, as we have shown mathematically, highly clustered representations can limit the dimensionality of representations, even when the positions of the clusters are random. Thus, to achieve the high dimensionality, the neuronal responses need to be diverse enough, which often means that they exhibit mixed selectivity [7, 8, 37] Our analysis of the response profiles revealed that they are indeed very diverse, as previously observed in other studies (see e.g. [1, 6, 12, 13]): even when there is some significant clustering, there is always the possibility that there is some additional structure within each cluster, given the cluster size. This diversity of responses within each cluster can greatly contribute to increasing separability, and hence, this diversity is potentially computationally important. Notice that the absence of clustering does not necessarily mean that the neuronal responses are entirely random. For example, different variables could be encoded with varying strengths, thereby elongating a non-clustered Gaussian distribution of responses along certain directions. Neurons with high selectivity to some variables could be considered ‘specialized’, but they would still be part of a continuous distribution of responses.

### Modularity and cell types

In the brain, there are several different neuronal types [38], highlighting a biological complexity that is especially marked for inhibitory neurons [39]. It is not unreasonable to expect that these different types of neurons would cluster in the response profile space. However, our analysis shows that only primary sensory areas are clustered, and even in these areas, the clusters are not well separated. One possible explanation is that the discrete nature of neuronal type is not directly reflected in their functional response to task variables. Another possibility is that the response profile reflects neuronal types that are not discrete or well separated. A recent article from the Harris-Carandini lab [40] analyzed the transcriptomic profile of neurons in the visual cortex and found a relation between the position of each neuron along a “transcriptomic axis” (found as the principal component in the high-dimensional space of 72 genes) and their activity modulation in different behavioral states of the subject. Crucially, while clustering in the transcriptomic space was found to correlate with putative cell types, this axis was defined as a “continuum,” hence, without clear-cut clusters. The transcriptomic space looks only partially clustered, with some different neuronal types arranged in continuous filaments. Further studies will be needed to assess whether other structures emerge in experiments involving more complex tasks (see also the last section of the Discussion). Future work will also explore the relationship between functional tuning (e.g., the RRR coefficients) and cell type embeddings estimated from spike waveform, autocorrelation [41], and eventually also transcriptomic/anatomical identity.

### Clustering of response profiles in other species

The recent article by the Kanwisher group [19] performed an analysis that systematically studies structures in the space of neuronal response profiles. In particular, they analyzed the visual and auditory systems of humans (fMRI) and monkeys (electrophysiology). Consistent with our results, when they examined mesoscale brain structures, they identified privileged neuronal axes that are preserved across individuals and reflect the large-scale organization of the brain. Their result is compatible with our observation in Fig. 3e-h, which shows clear clustering in the case of mesoscale sensory brain structures. Instead, when they repeated the analysis on a more local scale, within category-selective regions of the high-level visual cortex, they did not find the same structure, and they reported a distribution of the response profiles that would be comparable to the ones of our null models (these results are preliminary and mentioned in their Discussion). As the authors note, it is possible that this is a limitation of the resolution of fMRI, and that interesting structures emerge in electrophysiological recordings.

### Different measures of dimensionality

Measuring dimensionality in the presence of noise and determining whether it is high or low can be quite challenging [7]. This is why, in neuroscience, multiple methods are used to assess dimensionality. Each approach is different, and since dimensionality is expressed as a single number, it inevitably emphasizes only specific aspects of the representational geometry.

In our article, we always refer to the embedding dimensionality of the set of points representing different experimental conditions in the activity space[18]. These points represent patterns of activity recorded at the same time (not trajectories). To discount the dimensions due to the noise, we computed the Representation dimensionality of the average positions of the conditions. An alternative approach is to consider a cross-validated measure of Representation dimensionality [42]. Notice that separability and other measures related to it (see e.g. [6]), which consider the computational consequences of high dimensionality, are typically insensitive to noise, as they are also cross-validated measures.

Other recent works have introduced dimensionality measures that depend on the spatial and temporal scales considered [43]. For large scales, they get an estimate of the embedding dimensionality, and for short scales, the intrinsic dimensionality. While these quantities can reveal many other interesting aspects of the representational geometry, here we focused only on the embedding dimensionality because it is the one most relevant to the performance of a linear readout.

### High separability and other measures of performance

In all the brain areas that we analyzed, the representations are maximally separable when we consider only the independent conditions. In other words, when all pairs of conditions are separable, then all possible dichotomies (arbitrary ways of separating all the conditions into two groups with a hyperplane) are decodable by a linear readout above chance level. This serves as a critical warning for analyzing neural data: when independent conditions are considered, and dimensionality is sufficiently high, all variables associated with potential dichotomies can be decoded better than chance in all areas. Consequently, the decodability of a specific variable often lacks significance, as all other variables are also likely to be decodable.

In our data, we observed high separability in all the areas. This does not imply a complete absence of structure. First, because, as we showed, other measures of performance (e.g., the average separability) are more graded and vary across brain areas. Second, because representations can have the generalization properties of low-dimensional disentangled representations and still exhibit maximal separability [14] (notice that in this article both separability and the average decodability are close to 100%, so it is a more extreme situation than what we observed here). So, a learning process can lead to other interesting computational properties typically associated with low-dimensional representations without compromising the flexibility conferred by high separability.

It is also important to remember that separability is only one possible performance measure that is related to the dimensionality of the representations. A single number cannot summarize all important aspects of the dependence of the performance on the geometry, which in turn is determined by the diversity of the neuronal responses. This is why in the past, we considered also other measures of separability: the shattering dimensionality (here called ‘average separability’) introduced in [14] considers not only whether a dichotomy is linearly decodable or not, but also with what performance it is decodable. This is important as the chance level for decoding accuracy is often relatively low, and there is a wide range of accuracies above chance. As shown in the results, this quantity is also clearly related to the diversity of the responses, both in simulations and in the data.

A different approach, adopted in [6], considered a dichotomy that was decodable only if the performance exceeded a high threshold (85%) in the limit for large *N* (number of neurons) and large *p* (number of conditions). In this limit, the exact value of the threshold is actually less important, as predicted by Cover’s theorem (see also Suppl. material of [6]). However, estimating this limit is laborious, and hence we decided to use a different approach here, given the scale of the data and the higher level of noise compared to monkey recordings.

### The computational advantages of high separability

In feed-forward multi-layer artificial neural networks, it is important to have highly separable representations in the last layer because otherwise the number of input-output functions that the output units can implement could be severely restricted. However, this is not a strict requirement for intermediate layers.

So why are the neural representations so separable in the mouse cortex? One possible computational reason is related to the strong recurrent connectivity of the cortex. In artificial recurrent neural networks (RNNs), many computational properties, such as separability, depend on the dimensionality of the representations. One famous example is the memory capacity of the Hopfield network [44], which is rather limited for low-dimensional memories [15, 45]. Moreover, any RNN that needs to transition to a new state that depends on both the previous state and the external input requires neurons to non-linearly mix these two sources of information, which otherwise would constitute a concatenated input/recurrent state representation that is typically low-dimensional [8, 31].

Fortunately, increasing the dimensionality is relatively easy as non-linear random projections already do a surprisingly good job, both in RNNs [31, 46–48] and feed-forward networks [29, 49, 50]. More generally, if a network is initialized with random connectivity and the parameters are properly tuned, it is likely that the representations are already highly separable. Learning has numerous fundamental computational benefits, and it can certainly improve generalization and robustness to noise, as the network can extract the most relevant features for the task [51–54]. However, starting from high-dimensional representations is not that difficult, and networks operating in a lazy regime (i.e. a regime in which only the readout is trained and features are not learned) can already perform several tasks with a reasonable performance [51].

### The computational advantages of clustered response profiles

The brain is highly organized into functional and anatomical structures, which can be considered “modules” of the brain. This large-scale organization is well known; it has computational implications, and its emergence can be explained using general computational principles, at least in the visual system [55–57]. However, when one looks inside a particular brain area, the picture is less clear, though in several cases, it is possible to identify some form of modularity, which would lead to clustering in the response profile space. Two main forms of modularity have been studied in theory and experiments. The first modularity is observed for specific forms of disentangled representations that are aligned to the neuronal axes. Disentangled abstract representations are widely observed in the brain [14, 15, 17, 58] and are important for generalization. However, this geometry is compatible with both highly diverse neuronal responses (e.g., in the case of linear mixed selectivity) and with more organized categorical representations with modular structures in the neuronal response profile space. In the second case, each relevant variable would be represented by a segregated population of neurons (module), which would be a cluster in the selectivity space. This specific type of modular representation has been shown to be computationally equivalent to non-modular ones (although more challenging to compress under biological constraints [59]) but more efficient in terms of energy consumption and the number of required connections, at least under some assumptions [19, 60].

A second type of modularity has been characterized in artificial neural networks that are trained to perform multiple tasks [61, 62] or operate in different contexts [9, 63, 64]. In this case, it is possible to observe modularity, even when no additional constraints are imposed on metabolic costs and the number of connections [64]. In particular, two subtypes of modularity are considered: 1) *explicit* modularity, for which there are segregated populations of neurons that are activated in different contexts. In the activity space, this would imply that the conditions of different contexts are represented in orthogonal subspaces. Interestingly, the same geometry would be observed in the response profile space, and these representations exhibit clustering (i.e., they are categorical). 2) *implicit* modularity, for which the geometry in the activity space is the same as in the case of explicit modularity (it is a rotated version of the explicit modularity case), but in the response space, it is difficult to say whether clustering would be observed or not.

All these different types of modular representations share similar computational properties and allow us to generalize more easily, learn new structures more rapidly [64], and even enable some form of compositionality, in which the dynamics of sub-circuits can be reused in different tasks [62].

### Limitations of our study

Given all these computational advantages of modular structures, and the fact that categorical representations have been observed in some experiments [2, 10, 11], it is surprising that we observe significant clustering only in a few sensory areas. One possible explanation is that the assumptions made in the theoretical studies are incorrect and that the biological brain operates in different regimes. For example, the metabolic advantage of modular representation could be too modest compared to the enormous baseline consumption [65].

However, there are also other possible explanations. We looked for clustering using three ways of characterizing the response profile of each neuron (conditions space, regression coefficients of RRR, and time-averaged regression coefficients), two different algorithms for finding clusters (k-means and Leiden algorithm), and two statistical tests (one based on silhouette score, and one on ePAIRS). It is possible that other approaches will reveal some form of clustering or entirely different interesting structures in the statistics of the response profiles [66]. Moreover, we focused on the neuronal responses to the stimuli, and we did not consider the statistical properties of the spontaneous activity. Recent work shows that there is a relation between spontaneous activity patterns and the intra-PFC hierarchy[67], whereas our analysis of the response profiles did not reveal any categorical structure within the prefrontal module.

Finally, perhaps the most notable limitation of our study is that the IBL task is relatively simple, and the recorded animals are trained (in this case, often over-trained) to perform a single task. It is possible that repeating our analysis on a dataset that involves multiple tasks would actually reveal more clustering and specific modular structures. Indeed, Dubreil, Valente, et al. [63] showed that, in RNNs, simple tasks do not require clusters, whereas complex tasks do. Similar considerations apply to other theoretical studies [62, 64]. Future studies on multiple complex tasks will reveal whether the organizational principles that we identified are more general and valid in experiments that are closer to real-world situations.

## Acknowledgments

We are grateful to Fabio Stefanini, Mattia Rigotti, Marcus Benna, Ramon Nogueira, and Genevera Allen for numerous discussions about categorical representations and their analysis, to Tatiana Engel for the deep exchange of perspectives on dimensionality and clustering, and to our colleagues in the IBL for many fruitful discussions. We thank Yizi Zhang and Ji Xia for a code review of the RRR encoding model. LP is thankful to Jeff Johnston and James Whittington for their help in co-organizing the workshop “Are Neurons Interpretable?” (COSYNE, 2023), which played a big role in inspiring this work.

## Data Availability

The data used for this work is publicly available through the IBL and Allen Institute resources. The code used for the analysis is available on GitHub: github.com/lposani/decodanda (decoding and dimensionality analyses), https://github.com/realwsq/clustering-analysis for clustering analysis, and https://github.com/realwsq/brainwide-RRR-encoding-model (reduced-rank regression model).

## Methods

### Data structure

We used the International Brain Laboratory (IBL) public data release [4]. For each experimental session (178 in total), we collected time-series data on task, behavior, and electrophysiological recordings. These were segmented into trials based on key task events. The task recordings collected for each trial included information on the block prior as well as the stimulus contrast and location. The behavior recordings for each trial comprised the choice made, the outcome/ reward received, and the time-varying movement such as wheel movement velocity, whisker motion energy, and licks. Other behavior movements, such as paw movement, body motion energy, and pupil diameter traces, could be potentially included. However, we did not include them due to missing values in many sessions. The electrophysiological recordings for each trial contained time-varying spike trains of recorded neurons. All these recordings can be accessed directly via the IBL’s open API. The following section details the steps for preprocessing these raw data into data matrices for the encoding model.

#### Criteria for session inclusion

We iterated over all cortical regions and downloaded the related sessions. Sessions were included as long as all the behavior recordings (wheel velocity, whisker motion energy and licks) and electrophysiological data were in place. Some analyses required additional inclusion criteria, such as a minimum number of trials per condition. These analysis-specific criteria are discussed in the relevant sections below.

#### Criteria for trial inclusion

All trials from the left or right unbalanced blocks were included except when the animals did not respond to the stimulus in time (first movement time was longer than 0.8 s). Trials from the “50-50” balanced block were excluded from the analysis to avoid possible time artifacts arising from the fact that all these trials were exclusively recorded in the first 90 trials of the session.

#### Criteria for neuron inclusion

All neurons were included in the downloaded data as long as their mean firing rate was larger than 0.5 Hz and smaller than 50 Hz. For the selectivity and geometry analyses, we included only neurons that were predicted above a minimal threshold of min Δ*R*^2^ using the RRR model described below. Unless specified differently, we used min Δ*R*^2^ = 0.015. This threshold was necessary to avoid the confounding effects of neurons that do not encode any relevant variable, including those neurons that were recorded with a low signal-to-noise ratio. But the exact value of the threshold is not crucial for obtaining our results.

### Reduced-rank regression encoding model

In this section, we describe the reduced-rank regression (RRR) model used to analyze the selectivity profiles of single neurons. We start by describing the input variables and the target variables of the model, followed by the description of the model itself and its fitting procedure. Finally, we introduce a few quantities resulting from the fitted model that are key to the follow-up analysis. The notation that will be used are summarized in Table 1. The code for implementing and fitting the encoding model is available at https://github.com/realwsq/brainwide-RRR-encoding-model.

**Table 1.**
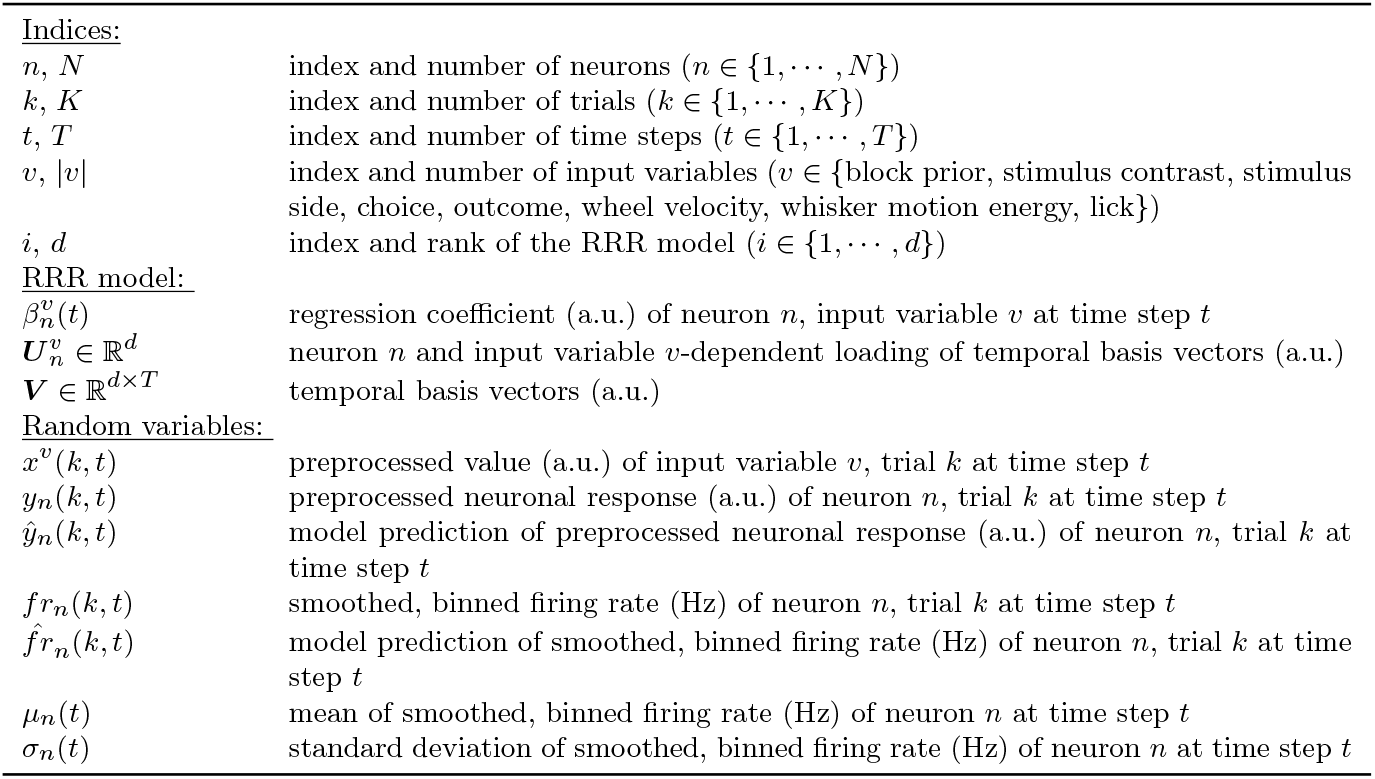
Notation.

### Input and target variables

#### Target variables

For each trial, we used the spike trains of the time window −0.2 to 0.8 seconds relative to stimulus onset, as the first movement time is typically less than 0.8 seconds [4]. The activity of each neuron was first binned at 0.01 s, divided by the size of the time bin, and then smoothed with a Gaussian filter with a standard deviation of 0.02 s. We tried to apply the linear time warping technique [68] so that the stimulus onset time and first movement or response time aligned across trials, but the results did not differ substantially. The resulting activity of neuron *n*, denoted as *fr*_*n*_, was organized into a matrix of shape *K*_*n*_ ×*T*, where *K*_*n*_ is the number of trials and *T* = 100 is the number of time steps per trial. *K*_*n*_ depends on *n* as neurons may have different numbers of trials if they were recorded in different sessions. Finally, for each neuron and each time step, we Z-scored the activity *fr* across trials to obtain the target variable *y* as follows (Suppl. Fig. 1b-d):

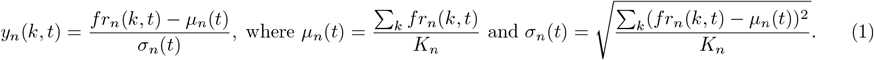

Since the squared error between the preprocessed data and model predictions was used in the loss function for optimizing the model (Sec 4), the applied normalization prevented biases due to inherent differences in activity scales and ensured that the predictions for all the neurons and time steps were optimized equally. Notably, the Z-score transformation is easily invertible, allowing the model’s predictions to be mapped back to the original units of firing rate, thus preserving interpretability.

#### Input variables

The input variables (*x*, Suppl. Fig. 1a) we considered can be divided into two types: discrete task-based variables and continuous movement variables. Discrete task-based variables include task-related features, such as the block prior, stimulus contrast, stimulus side, choice, and outcome. These are listed below:

- **Block**: The prior probability for the stimulus to appear on the left side is either *p*(left) = 0.2 (right block), or *p*(left) = 0.8 (left block). We used one input variable to encode the block prior: −1. representing *p*(left) = 0.2, and +1. representing *p*(left) = 0.8. As noted above, we excluded trials from *p*(left) = 0.5 unbiased block.
- **Contrast**: The stimulus contrast is 0%, 6.25%, 12.5%, 25%, or 100%. One variable was used to encode the stimulus contrast: 0. representing 0% contrast, 1. representing the low contrast (≤ 12.5%), and 4. representing the high contrast (> 12.5%).
- **Stimulus**: The stimulus location is either on the left side (+1.) or the right side (−1.).
- **Choice**: The choice is indicated by the turning of the wheel: clockwise (+1.) or counterclockwise (−1.).
- **Outcome**: The outcome is either a water reward (+1.), or negative feedback (−1.).

Since these values are static, all the time points share the same values (e.g., see Suppl. Fig. 1a).

Continuous movement variables included both instructed (e.g., licking and wheel velocity) and uninstructed (e.g., whisker motion energy) movement. These are listed below.

- **Wheel**: The velocity of the wheel movement (radian per second) per time bin.
- **Whisking**: The whisker motion energy per time bin is calculated as the motion energy for a square of the left/ right camera roughly covering the whisker pad. The maximum value between the left and right whisker motion energy was used.
- **Lick**: The number of licks per time bin.

A few preprocessing steps were applied separately to each movement variable: for each trial, we first read out the continuous behavior of the time window −0.2 to 0.8 seconds relative to stimulus onset and interpolated into 0.01 s time bins. Then, to account for the activity that was shifted in time, for each session, we computed the mean time-lagged correlation between the neuronal activity and the movement traces averaged across neurons and trials and shifted the movement traces so that the zero-lagged correlation was maximized. Lastly, significant differences were observed in the variance of input values across trials, both for different input variables and different time steps. To ensure optimal performance and clarity in interpretation, we Z-scored the values for each input variable and time step across trials in the same way as Eq 1. Examples of the resulting input variables are shown in Suppl. Fig. 1a.

### The reduced-rank regression model

#### Linear encoding model

For each neuron *n*, we describe its temporal responses as a linear, time-dependent combination of input variables (Suppl. Fig. 1ab):

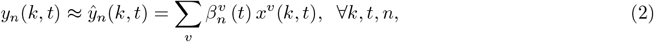

where

- *y*_*n*_(*k, t*) is the preprocessed neuronal activity of the trial *k* ∈ {1, · · ·, *K*_*n*_} and time step t ∈ {1, · · ·, *T* }.
- *ŷ*_*n*_(*k, t*) is the corresponding model prediction, given by the value of the equation on the right-hand side.
- *v* represents the relevant input variables included in the model. *x*^*v*^(*k, t*) is the preprocessed value of the input variable v for the trial k and time step *t*.
- 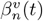 is the effect size of the input variable *v* at time step *t*. It is further referred to as the regression *coefficient*.

#### Low-rank coefficient matrix

The time-varying coefficients 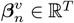 are the weighted sum of a set of temporal basis vectors shared across all the neurons and input variables, that is

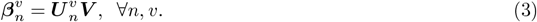

Here, 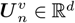 is the neuron *n* and input variable *v*-dependent loading of *temporal basis vectors* ***V*** ∈ ℝ^*d×T*^ . Specifically, we considered sharing a single set of temporal basis vectors across all the neurons from all the brain regions and across all the input variables. We verified that this restriction did not compromise the goodness-of-fit. The rank *d* is generally a value much smaller than the number of time steps *T* . See Suppl. Fig. 1e for an example decomposition. Sharing the temporal bases across neurons and input variables significantly reduces the number of parameters (Suppl. Fig. 1f). Let *N* be the number of neurons, *T* be the number of time steps, and |*v*| be the number of input variables. An unconstrained full-rank coefficient matrix employs *N* ×|*v*| ×*T* parameters, while a reduced-rank coefficient matrix of the same shape only needs *N* × |*v*| × *d* + *d* × *T* parameters. Since *N, T* are typically much larger than *d*, the reduction in parameters is on the order of *T*, i.e., around 100-fold.

### Comparison to previous regression models

Well-established linear encoding models include generalized linear model (GLM) [4, 20] and kernel regression model [21]. Below, we first describe these two models and then discuss how they compare to our RRR model. GLM is expressed as

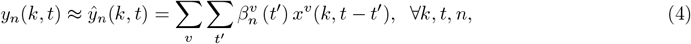

where the input filter vector, 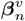, composed of 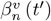 over a neighboring time window, is factorized as

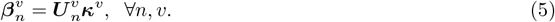

Here, *κ*^*v*^ represents pre-specified temporal basis vectors, which are not trained. Adapted from [4] ^1^, we used the same set of raised cosine “bump” functions in log space for Suppl. Fig. 3b. The kernel regression model employs the same linear formulation as the GLM (Eq 4). However, instead of relying on pre-defined temporal basis vectors, it allows these vectors to be trainable:

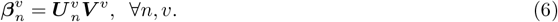

Both the kernel regression model and our RRR model fall into the category of reduced-rank regression models. The primary distinction lies in the set of input features used by each approach.

In our RRR model, the influence of input variables on neural responses, 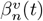 can vary over time. In contrast, both the GLM and the kernel regression model assume that the influence is time-independent. Therefore, for example, whether the subject response time is early or late makes no difference and will modulate the neural responses the same way. Instead, these models provide a more descriptive account of input effects by allowing neural responses to depend on inputs from neighboring time steps. While this feature is absent in our current RRR model (Eq. 2), it could be incorporated in future extensions.

### Estimation of the parameters

The parameters of the RRR model include a shared temporal bases matrix ***V*** of size *d*×*T* and loading vectors 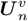 of length *d* for each input variable *v* and neuron *n*. The approach we adapted to fit the parameters was to minimize the ridge-penalized mean square loss:

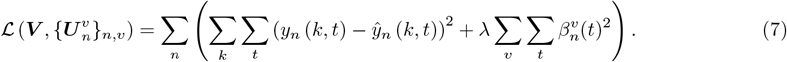

Minimizing this particular loss function is straightforward as a closed-form solution exists [69]. In practice, we chose to use the L-BFGS optimization algorithm to compute the optimum.

Moreover, to optimize the model hyperparameters, namely the rank *d* and the regularization penalty *λ*, we implemented a 3-fold cross-validation technique across trials. First, the dataset was stratified based on a composite target label that included the block prior and stimulus contrast to ensure each fold was representative of the entire dataset. Then, for each combination of *d* and *λ*, the dataset was partitioned into three subsets by trials, utilizing each subset in turn for testing the model while the remaining data served as the training set. Finally, the combination of *d* and *λ* that yielded the lowest average test error across all folds was selected. *d* = 5 turned out to be the optimal number of temporal bases (Suppl. Fig. 1e).

### Estimating the goodness of fit

We used the 3-fold cross-validated *R*^2^ to measure the goodness-of-fit of single-trial predictions. For each session’s data, we randomly sampled one-third of the trials as the test set held out during training. Once the model is trained - using the remaining two-thirds of trials - we computed the *R*^2^ between the model predictions 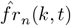 ^2^ and the actual neuronal activity *fr*_*n*_(*k, t*) on the test set. We repeated the whole split-train-test process three times and computed the mean of the three cross-validated *R*^2^ as the measure of goodness-of-fit.

#### Selectively modulated neurons

Conceptually, we distinguish two types of task modulation: the *selective modulation* and the *non-selective modulation*. Selective modulation, captured by 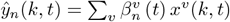 is induced by the input variables and varied trial-by-trial. Non-selective modulation, captured by the mean time-varying response 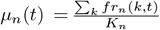 is locked to the key events of the trial (stimulus onset in this case) and does not vary trial-by-trial. Both types play a significant role in modulating neuronal responses. See Suppl. Fig. 3 for example selectively modulated and non-selectively modulated neurons. In this work, we focus mostly on the neuronal responses selectively modulated by the task. To distinguish the variation explained by the selective modulation from the non-selective modulation, we use the trial-average estimate as the null model that does not consider any effect of the input variables:

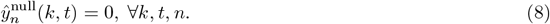

The outperformance, Δ*R*^2^, defined as:

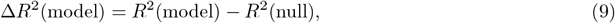

captures the overall selective modulation of all the input variables combined.

Only selectively modulated neurons, identified as Δ*R*^2^(RRR) ≥ 0.015 (Suppl. Fig. 3a), are included in the further analysis.

### Computing the selectivity profiles of single neurons to the input variable

To compute the selectivity profiles of single neurons to the individual variables, we used the estimated coefficient 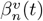. For the clustering analysis of Fig. 3, Suppl. Fig. 10), we took the sum of the coefficients across time as a measure of the total selectivity of neuron *n* to input variable *v*.

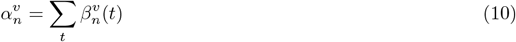

Note that by normalizing the neuronal responses and input variables in the preprocessing steps, we ensured that the unit-free coefficients 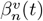 are not affected by the neuron’s mean firing rate or the inherently different scales in different input variables and can be compared directly across neurons, input variables, and time steps. Thus, 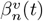 can be interpreted as the expected change in normalized neuronal activity *y*_*n*_ per one standard deviation change in the input variable *x*^*v*^ at time *t*, and 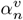 can hence be thought as the expected total change across the whole trial. Also, note that the sign of 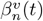 typically does not change over time *t*. See Suppl. Fig. 1a for examples.

The selectivity 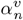 captures whether individual neurons are selectively modulated by the given variable *v* or not (see Suppl. Fig. 2 for examples of strongly selective neurons). In the selectivity analysis (Fig. 2ef), when the goal is to estimate the absolute modulation of an input variable, 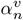 is calculated as 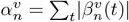.

### Computing the autocorrelation timescale of neural responses to task variables

Given a matrix of single-neuron responses to task variables ***ŷ***_*n*_ ∈ ℝ^*K×T*^ with *K* being the number of trials and *T* the number of time steps per trial, we can compute the corresponding autocorrelation timescale. To compute the timescale, we first calculate the time-lagged unnormalized autocorrelation sequence

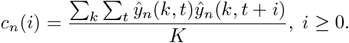

We then linearly interpolate the autocorrelation sequence so that *c*_*n*_(*i*) is spaced at 1 ms resolution (original 10 ms). The timescale *τ*_*n*_ is approximated by the time the sequence first reaches half its peak value (i.e., *c*_*n*_(0)). The timescale of brain area *a* (Fig. 2d) is further determined by averaging the values over all the selectively modulated neurons within this area.

### Clustering Analysis

To test for the presence of functional clusters, we followed the steps explained below. The required inputs include each relevant neuron’s response profile and original session ID. Two types of response profiles can be considered: the estimated selectivity to individual input variables (Eq 10, Fig. 3, Suppl. Fig. 10, referred to as clustering analysis in the variable selectivity space) or the average response in each experimental condition (Suppl. Fig. 9, Suppl. Fig. 10, referred to as clustering analysis in the conditions space). Performing clustering analysis in the selectivity space arguably has a few advantages:

- It reduces the dimensionality in an interpretable and informed way. If we have |*v*| variables, then there are at least 2^|*v*|^ conditions, assuming all the variables are discrete and have more than one different value.
- It mitigates the issue of unbalanced or even missing conditions.
- It reduces the noise in the estimation of the response profile. As shown in Suppl. Figure 3, neural responses are very noisy, and simple averaging may be non-satisfactory (Figure 2b). The selectivity estimated from the encoding model, in comparison, provides a more reliable account of the task-driven variance in the neural responses.

The code for the clustering analysis is available at https://github.com/realwsq/clustering-analysis.

#### Clustering analysis in the variable selectivity space

We summarize our clustering pipeline below (see also Fig. 3a for schematic illustration).

1. Check whether there are more than 50 neurons and only continue if so.

2. Given the selectivity profile of each neuron, run k-means clustering algorithm (with 100 random initializations) with the number of clusters *k* varied from 3 to 20.

3. Then, select the optimal clustering result by maximizing the silhouette score. The Silhouette score is defined as 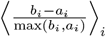 where *i* is the index of the neuron, 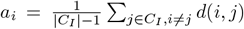 is the mean Euclidean distance intra-cluster and 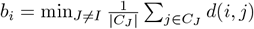 is the min Euclidean distance outside cluster (Fig. 3a).

4. Iterate over the resulting clusters and check whether there is a cluster whose total silhouette score summed over all neurons is mainly contributed by neurons from one single session (> 90%). If so, remove neurons from that cluster and session, and repeat steps 1-4.

5. Sample the same number of data points from the Gaussian distribution with the mean and covariance matrix matched to the data values and compute the sampled data’s null silhouette score following steps 2-3.

6. Repeat step 5 100 times and pool the null silhouette scores to form the null distribution. Finally, compute the z-score of the data silhouette score with respect to the null distribution.

#### Clustering analysis in the conditions space

When clustering in the space of mean firing rate, two additional preprocessing steps are required. First, we normalized each neuron’s mean firing rates separately across conditions to prevent clustering driven solely by overall firing rate differences between neurons. Second, since the number of conditions and the dimensionality of the activity profile is high, we reduced the dimensionality using principal component analysis. Additionally, we modified the null model by replacing the multivariate Gaussian distribution with a multivariate log-normal distribution to better capture the lower-bounded and heavy-tailed nature of mean firing rates. Neurons with zero activity were excluded from this analysis.

The clustering analysis in the conditions space follows these steps (see also Suppl. Fig. 9 for schematic illustration):

1. Check whether there are more than 40 neurons and only continue if so.
2. Z-score the mean firing rates for each neuron.
3. Reduce dimensionality using PCA, retaining components that capture 90% of the total variance.
4. Run k-means clustering algorithm (with 100 random initialization), varying the number of clusters *k* from 3 to 20.
5. Select the optimal clustering result by maximizing the silhouette score.
6. Iterate over the resulting clusters and check whether there is a cluster whose total silhouette score summed over all neurons is mainly contributed by neurons from one single session (> 90%). If so, remove neurons from that cluster and session, and repeat steps 1-5.
7. Sample the same number of data points from the multivariate log-normal distribution with the mean and covariance matrix matched to the log data values, then compute the sampled data’s null silhouette score following steps 2-5.
8. Repeat step 7 100 times and pool the null silhouette scores to form the null distribution. Finally, compute the z-score of the data silhouette score with respect to the null distribution.

#### Measuring the similarity between cluster and area labels

The similarity between functional clusters and anatomical area labels was quantified using the Rand Index (RI), which measures the agreement between two labelings of the same dataset. For each pair of neurons, agreement occurs if both are assigned to the same cluster in both labelings, or to different clusters in both. The RI is defined as the fraction of agreeing pairs among all possible pairs. To assess statistical significance, we computed a z-scored RI by comparing the observed RI to a null distribution obtained from 10,000 random shufflings of the area labels. A significantly elevated z-scored RI indicates that functional clustering aligns closely with anatomical organization. To avoid biases toward areas or modules with larger neuron numbers, we considered the 100 best-encoded neurons from each module or area in the analysis of Fig. 3e, f and Suppl. Fig. 11.

### Modified ePAIRS test

Given the average responses of single neurons in each experimental condition, we performed the ePAIRS test as follows (see also Suppl. Fig. 9 for schematic illustration):

1. Z-score the mean firing rates for each neuron.
2. Reduce dimensionality using PCA, retaining components that capture 90% of the total variance.
3. Calculate the cosine distance to its nearest neighbor for each neuron.
4. Calculate the empirical median of these nearest-neighbor distances as the aggregated nearest-neighbor angle.
5. Sample the same number of data points from the multivariate log-normal distribution with the mean and covariance matrix matched to the log data values, then compute the null nearest-neighbor angle of the sampled data following steps 1-4.
6. Repeat step 5 5000 times and pool the null nearest-neighbor angles to form the null distribution. Finally, the z-score of the data aggregated nearest-neighbor angle with respect to the null distribution is computed.

### *α*-diversity

To measure *α*-diversity, we took the participation ratio of the *N* × *V* matrix of alpha coefficients resulting from the RRR analysis described above. The PR quantifies the effective dimensionality of a set of data points by measuring how evenly the variance is distributed across the eigenvalues of its PCA decomposition [29]. The PR is defined as:

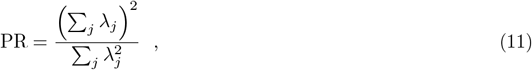

where *λ*_*j*_ is the j-th eigenvalue of the *N* × *N* covariance matrix. A higher PR indicates that the variance is more evenly spread across multiple dimensions, suggesting a higher effective dimensionality of the cloud of points in the high-dimensional space. Conversely, a lower PR implies that the variance is concentrated in fewer dimensions, indicating a lower effective dimensionality.

The number of neurons in individual areas was soft-equalized by taking a random subsample of *N*_0_ = 120 neurons when *N* > *N*_0_. In these cases, the participation ratio was computed over 100 random subsets, and the average was taken as the value of *α*-diversity.

### Analysis of population representations

#### Data Preparation

For the analysis of population neural representations, we used the same sessions as described above. For each trial within a session, we thus have a collection of *N*-dimensional population activity vectors **f** ^*t,k*^, where *k* ∈ {1, *P*} is the trial index within the session, and *t* ∈ {1, *T* } is the time-bin index within each trial. For the analysis below, we used data from 0 to 1000ms after the stimulus onset, so to capture a variety of sensory and behavioral variables. We then labeled each time bin according to the value of four binarized cognitive, sensory, and movement variables:

- **Block**: left (20-80) vs. right (80-20) prior block.
- **Contrast**: we binarized the contrast into low (0-0.125) vs. high (0.25-1.0) values.
- **Stimulus**: left vs. right side of the screen.
- **Whisking**: we binarized the whisking power using the distribution of whisking power values within each session. Time bins where the mouse was whisking with a power larger than the 50 percentile across the distribution were annotated as *high*, while those below the 50 percentile were annotated as *low*.

These variables were chosen so that they span movement, cognitive, and sensory variables while ensuring that all the *M* = 16 conditions (combinations of the four variables) were well represented in the data. For example, we could not add Choice as a variable since mice are overtrained in the task and, as a consequence, make very few mistakes when Block and Stimulus are aligned (e.g., choose “left” when the block and the stimulus are both “right”.)

For each condition *c* ∈ {1, *M*} (for example, Whisking = high, Contrast = low, Stimulus = left, Block = right), we first identified those trials where that specific combination of variables was present. We then defined a collection of “conditioned trial” population activity vectors {**f** ^*k,c*^} as the mean firing rate of the population of neurons conditioned to the specific condition in each trial. Given a trial *k* and a neuron index *i*, the mean firing rate was computed as

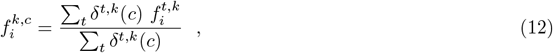

where *i* indicates the neuron index and *δ*^*t,k*^(*c*) = 1 if the time bin *t* in trial *k* corresponds to the condition *c*, and = 0 otherwise. These conditioned trial population vectors are the data samples that will be used for the dimensionality and decoding analyses below. Across all analyses, we considered only those recording sessions where each condition was present in at least *M*_*min*_ = 5 trials.

### Representation Dimensionality

To estimate the Representation Dimensionality of a neural geometry, we computed the PR of the *centroids* **f**_*c*_ of the *M* conditions, defined as the average activity pattern across all trials of the same condition:

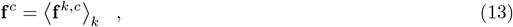

To compute the PR for the set of centroids, we first calculated their covariance matrix, normalizing each neuron’s mean activity vector by subtracting from each 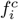 the mean across conditions *c* and dividing them by their standard deviation. We then performed Principal Component Analysis (PCA) on this covariance matrix to obtain its eigenvalues, *λ*_*j*_.

The number of neurons in individual areas was soft-equalized by taking a random subsample of *N*_0_ = 120 neurons when *N* > *N*_0_. In these cases, the participation ratio was computed over 100 random subsets, and the average was taken as the value of Representation Dimensionality.

### Cross-validated Decoding

We used Decodanda [70] (www.github.com/lposani/decodanda) to perform a cross-validated, class-balanced decoding analysis of different combination of condition labels from the neural activity within individual trials (condition trial vectors **f** ^*k,c*^). See individual sections below for additional detail on the data input structure of our decoding analyses. As a decoder, we used a scikit-learn SVM classifier with linear kernel [71]. To ensure that results were comparable across regions, which might have a different number of recorded neurons, we created a pseudo-population by resampling all the recorded neurons within each region to a fixed number *N* = 4000. Similarly, we re-sampled the same number of pseudo-population for each analysis (*T* = 100 patterns per condition). Note that simultaneously recorded neurons were always kept together during resampling, so as to keep the noise correlations intact within the pseudo population [70]. All cross-validated decoding analyses were performed using the following Decodanda parameters: `training_fraction` = 0.8, `cross_validations` = 100, `ndata` = 100.

### Finding the Independent Conditions

To find the number of independent conditions encoded in the activity of a population of neurons, we developed an iterative algorithm based on linear decoding. The algorithm followed the steps below, and is shown in action on one example region in Suppl. Fig. 5.

1. First, we performed a decoding analysis of the condition label *c* from trial population vectors **f** ^*k,c*^ using a set of binary linear classifiers: For each pair of conditions (*c*_*i*_, *c*_*j*_), we estimated a cross-validated decoding performance *φ*(*c*_*i*_, *c*_*j*_), resulting in an initial *M* × *M* condition-condition decoding matrix (C_0_, see Suppl. Fig. 5) defined as C_0_(*ij*) = *φ*(*c*_*i*_, *c*_*j*_).
2. We then chose a decoding threshold *φ*_min_ = 0.666; the pairs of conditions whose 1-vs-1 decoding performance was smaller than *φ*_min_ were defined as “dependent”. Using this threshold, we defined a binary *dependency* matrix D defined as D_0_(*i, j*) = 1 if *φ*(*c*_*i*_, *c*_*j*_) < *φ*_min_, and C_0_(*i, j*) = 0 otherwise.
3. Then, we used the Bron-Kerbosch algorithm [72] to find all the cliques, i.e., subgroups of fully-connected nodes, in the undirected graph defined by the dependency matrix D_0_. This process allows us to identify whether there are groups of conditions that are all non-decodable from each other (dark squares in the sorted matrix in Suppl. Fig. 5).
4. Next, we identified the largest clique and grouped together all the trials of the conditions within that group into a new, *merged* condition (see arrows and “merge” conditions in Suppl. Fig. 5).
5. We repeated steps 1 and 2 with the new reduced set of conditions, yielding a new C_*t*_ and a new D_*t*_ matrix of a different size *M*_*t*_; *t* denotes the iteration step.
6. We then repeated step 3 and 4, and iterated the whole process (1-4) until all the merged and remaining conditions were found to be independent, i.e, the dependency matrix 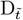 was diagonal. The number of independent conditions was then defined as the size of the final dependency matrix: 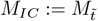.

### Separability and Average Decodability

Separability quantifies how many random dichotomies (equally-sized groups) of experimental conditions can be decoded from neural activity using cross-validated linear classifiers. To estimate the Separability of a neural population, we followed the following steps:

1. First, we randomly divided the set of *M* or *M*_*IC*_ independent conditions into two equally-sized groups (dichotomy).
2. Given the dichotomy *d*, we then measured the cross-validated decoding performance *φ*_*d*_ of a linear classifier trained to report whether individual condition trial vectors **f** ^*k,c*^ belonged to conditions within one or the other dichotomy groups. This decoding analysis was performed as described in the Decoding section above, resampling a fixed large number of neurons (*N* = 4000) and a fixed number of trials (*T* = 100) per condition for all regions to ensure performances could be compared across regions.
3. The random dichotomy assignment and decoding (steps 1, 2) was then repeated *n* = 200 times to obtain a set of decoding performances {*φ*_*d*_}.
4. the decoding analysis was then repeated for *n* = 200 times with shuffled condition labels across the population vectors to obtain a distribution of null decoding performance values {*φ*^null^}
5. Separability was then defined as the fraction of decodable dichotomies, i.e., the fraction of *φ*_*d*_ larger than the 99 percentile of the null population {*φ*^null^}. Average Decodability (AD) was defined as the mean decoding performance across the *n* = 200 random dichotomies.

The distributions of decoding performances for all the analyzed regions are shown in Suppl. Fig. 17.

### Synthetic data

#### Modeling uneven and categorical selectivity

We generated synthetic population responses for *N* neurons to all 2^*V*^ configurations of *V* binary variables *x*_*v*_ ∈ {−1, 1}. Each neuron’s response combined linear and quadratic selectivity,

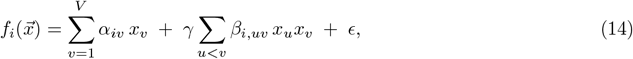

where *γ* controls the strength of nonlinear interactions and *ϵ*_*i*_ ∼ 𝒩 (0, σ) introduces trial-to-trial variability. For each condition 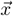, we generated *T* independent trials and used the data in the decoding analyses as performed for real spiking data.

#### Sampling the selectivity structure

The coefficients (*α*_*iv*_, *β*_*i,uv*_) were sampled in a feature space of dimension 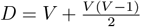, which groups all linear and quadratic features on an equal footing. To generate either *categorical* or *uneven* selectivity, we first drew *k* cluster centroids in the *D*-dimensional feature space and repeated each centroid *k*_*N*_ = *N*/*k* times, yielding *N* prototype vectors. These prototypes define the coarse structure of selectivity across neurons. To modulate within-cluster diversity, each prototype was perturbed by an additive diversity vector

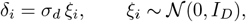

where *σ*_*d*_ = − log(*c*) is set by a *categorical specialization* parameter *c* ∈ (0, 1) (this is the x-axis in Suppl. Fig. 12b). Larger *c* produces more tightly clustered selectivity (strong categorical structure), whereas smaller *c* yields more dispersed coefficients. To model *uneven* selectivity, we introduced anisotropy across the *D* feature dimensions by using *k* = 1 (no clustering) and scaling the diversity terms by a geometric decay profile:

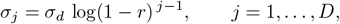

where *r* ∈ (0, 1) is the *global specialization* parameter. When *r* ≪ 1, all dimensions contribute equally; when *r* ≈ 1, variance is concentrated in a low-dimensional subspace, producing a strongly uneven selectivity spectrum. The final coefficients, including categorical and/or uneven structure, were

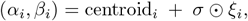

with the quadratic coefficients additionally scaled by *γ* to control nonlinearity.

#### Trial generation

For each of the 2^*V*^ binary stimuli, we computed 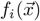 via the linear–quadratic form above, and generated *T* noisy samples per condition,

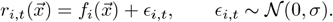

This procedure ensures that trial variability is independent across neurons and conditions, and that all structure in the population code arises exclusively from the geometry of the sampled coefficients.

#### Parameterizing categorical and uneven specialization

The two specialization parameters (*c, r*) thus independently control:

- the *categorical* structure of selectivity (number and separation of clusters, via *c* and *k*);
- the *unevenness* of selectivity across feature dimensions (anisotropy of coefficient variances, via *r*).

By sweeping either *c* (categorical specialization) or *r* (global specialization) while keeping the other fixed, we isolated the effects of clustered versus uneven selectivity on representational dimensionality, separability, and independent condition structure. For the analyses in Suppl. Fig. 12, we used *N* = 100 neurons, *V* = 3 variables (for a total of *P* = 8 conditions), *T* = 20 samples per condition, and *γ* = 0.25.

### Exploring the relation between dimensionality and separability

To analyze how separability changes with the dimensionality of the geometry in the activity space, we performed a series of synthetic explorations shown in Suppl. Fig. 14. In these simulations, P centroids are randomly sampled from a Gaussian distribution spanning an L-dimensional subspace of the N-dimensional activity space. Each centroid vector is normalized. Trial-to-trial variability is then added to the centroids with a standard deviation *σ*, scaled with 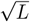 to keep the signal-to-noise level constant when pairwise distances between centroids increase with L. This obtains a TxN activity matrix for each of the P conditions. This synthetic activity is then analyzed with the same pipeline used for the cortical data, yielding values of Representation dimensionality, Separability, and Average Decodability shown in Fig. 14d, where we used P=16, N=100. For simulations in Fig. 14f, we fixed L=14 and stretched multiplied the first dimension of each centroid of a factor *γ* to stretch the geometry along a single axis.

### Theoretical considerations on the relationship between dimensionality and clustering

#### The conditions space and the neural space have the same dimensionality

As explained in Fig. 1a, response profiles of single neurons can be thought of as rows of a matrix *X* whose columns define the geometry of conditions in the neural space. The PR in the conditions space is computed from the eigenvalue spectrum of the covariance matrix of the rows of *X*, i.e., *XX*^*T*^ (assuming zero mean), while the PR in the activity space is computed from the eigenvalues of the covariance matrix of the columns of *X*, i.e., *X*^*T*^ *X*. Given a matrix, the eigenvalues of its covariance matrix are the squared singular values of *X*. Let *X* = *USV* ^*T*^ be the SVD decomposition of *X*, where *S* is the diagonal matrix with singular values on the diagonal, then *X*^*T*^ = *V S*^*T*^ *U* ^*T*^ . As *S* = *S*^*T*^, the eigenvalue spectrum of *X*^*T*^ *X* and *XX*^*T*^ is the same. Thus, the participation ratio of the conditions space is the same as that in the neural space.

#### Mathematical derivation of the PR of Gaussian clusters

We consider a data model with *N* features (neurons) and *M* observations (conditions), in which observations are sampled i.i.d. as

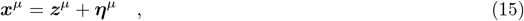

where ***z***^*µ*^ and ***η***^*µ*^ are both vectors in ℝ^*N*^ and represent the clustered and heterogeneous part of the data, respectively. More precisely, ***z***^*µ*^ is sampled from a normal distribution 𝒩 (0, **B**) that has a clustered covariance matrix, i.e. *B*_*ij*_ = 1 if *i* and *j* belong to the same cluster and *B*_*ij*_ = 0 otherwise. We call *k* the number of clusters and assume that all clusters have the same number of neurons *N*_*c*_ = *N*/*k*. In contrast, the heterogenous part ***η***^*µ*^ is sampled from 𝒩 0, *σ*^2^**I**, where **I** is the identity matrix. Our goal is to compute the participation ratio (PR) of this representation, which we define as

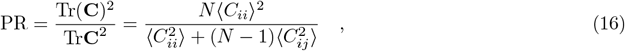

where the averages are across neurons and the matrix **C** is the *sample* neuron-by-neuron covariance matrix, i.e. 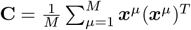. We note that this definition assumes that the sample mean of both ***z*** and ***η*** are negligible or have been subtracted.

We are interested in the regime in which *N* → ∞ while *M* is allowed to be small, as it often happens in controlled experiments. Small *M* might cause the sample covariance matrix to differ substantially from the true covariance matrix. Defining **C**^*c*^, **C**^*h*^, and **C**^*ch*^ as the sample covariance matrices of ***z, η***, and the cross-covariance between ***z*** and ***η*** respectively, we have that

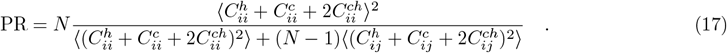

Therefore, we need to evaluate the first and second moments of both diagonal and off-diagonal elements of all covariance and cross-covariance matrices. The diagonal elements of these matrices have the following statistics:

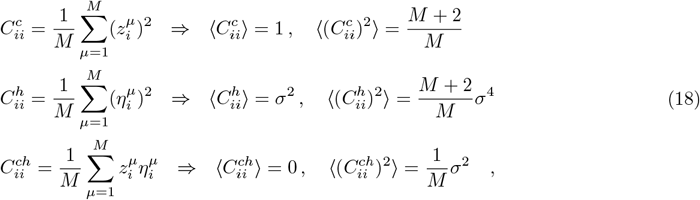

and

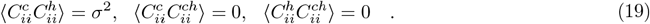

For the off-diagonal elements, we have

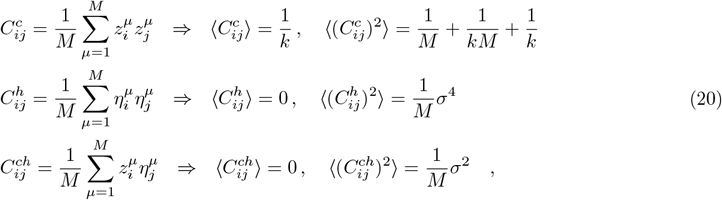

And

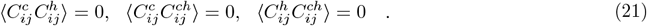

Most of the expressions above can be straightforwardly derived by writing down the definition of the sample covariance matrix for a zero-mean variable and then performing the average over neurons. To illustrate this procedure, let us consider one of the most involved terms:

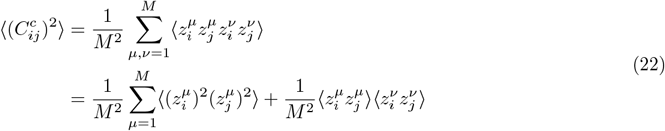

The probability that *z*_*i*_ and *z*_*j*_ are part of the same cluster is given, for large *N*, by 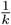 . The expression for 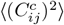 then becomes

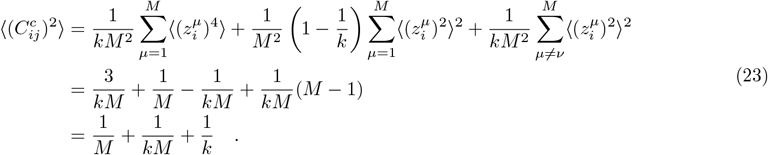

The other terms can be computed following the same steps.

Given that *k, M* are finite, we can approximate the PR for large *N* as:

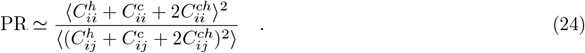

Expanding the square and using the results above for the first and second moments of the covariance matrices, we get to our final expression:

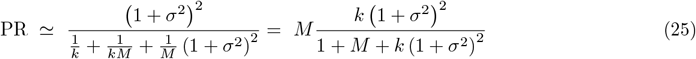

From the mathematical expression in Eq. **??**, we can see that, in the limit of perfect clusters (*σ* → 0), the function is either limited by the number of rows-neurons (in this case, the *k* perfect clusters) or columns-conditions *M*, coherently with the intuition above:

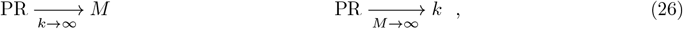

However, things become more nuanced when *k, M*, and *σ* are finite and non-zero. First, as shown in Fig. 5e, when *k* and *M* are kept fixed, the Representation dimensionality decreases with the clustering quality (expressed as the average silhouette score of a population of Gaussian clusters with given *k, M*, and *σ*). Second, if we fix the quality of clusters and the number of conditions (Fig. 5e, left), we see that dimensionality increases with the number of clusters, with a magnitude that is larger for high silhouette scores (categorical representations). This behavior is complementary to a recent work that showed the inverse relation, i.e., that constraining the dynamics generated by recurrent neural networks to a low dimensional manifold implies a small number of functional types (categorical clusters) [73]. Finally, when fixing the number and quality of clusters, the dimensionality is determined by the number of conditions, with a magnitude that is larger for low silhouette scores (non-categorical representations, Fig. 5e, right).

## Extended Data

**Supplementary Figure 1.**
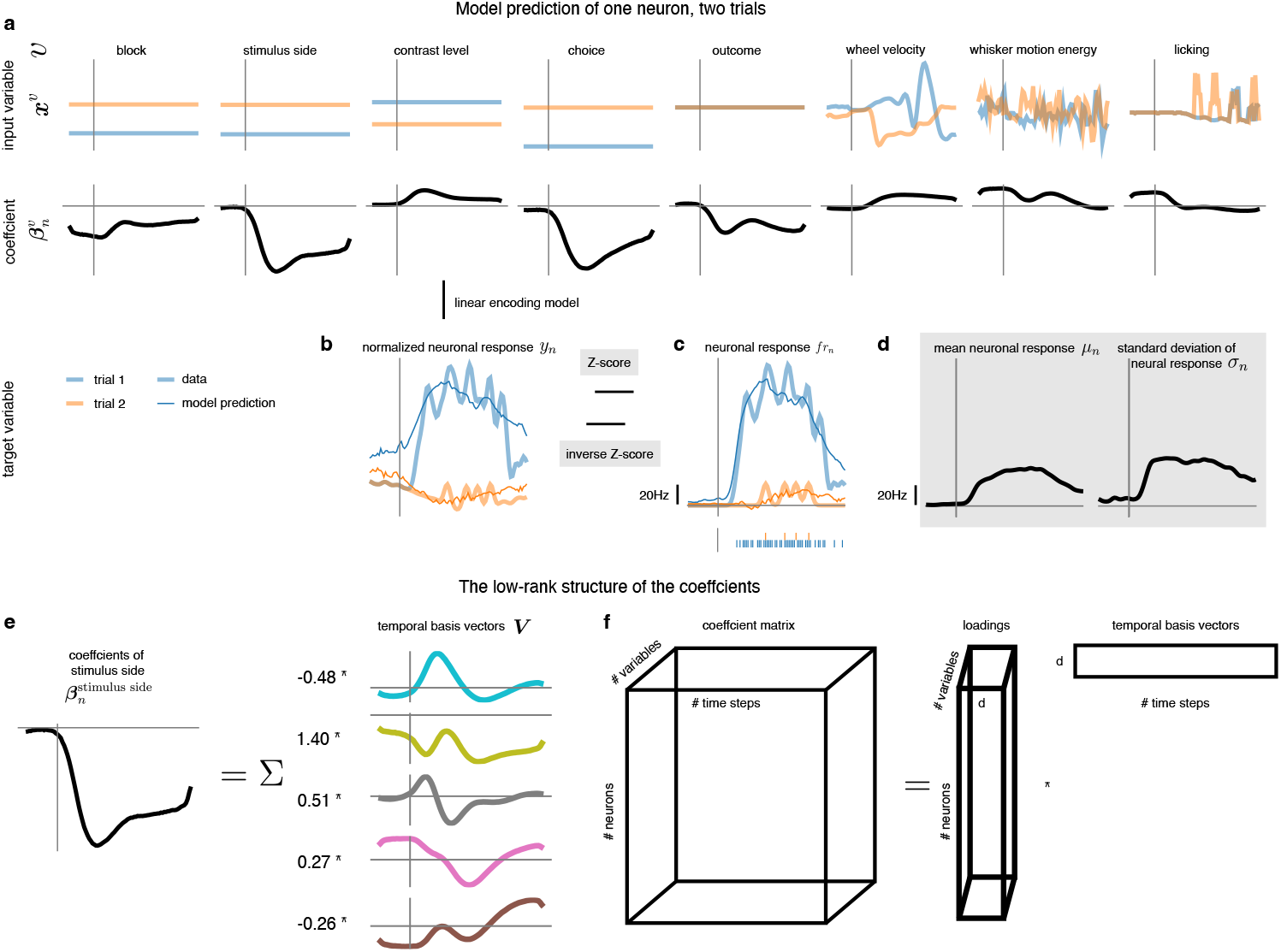
Visualization of the RRR encoding model. **a-d** Illustration of how the RRR encoding model predicts the neuronal responses using two example trials (blue and orange lines). Importantly, the model coefficients are time-dependent (a), and the inputs (*x*) and outputs (*y*) of the RRR encoding model are normalized across trials (b-d, Methods). The prediction (thin and opaque lines) overlays the data (thick and transparent lines). The gray vertical lines indicate the onset of the stimulus (0.2 s), and the gray horizontal lines indicate the zero value. **e-f** Coefficients are enforced to be weighted sums of a small set of temporal basis vectors. **e** Example decomposition of the stimulus side coefficients in (a). **f** Schematic plot showing the low-rank structure of the coefficient matrix.

**Supplementary Figure 2.**
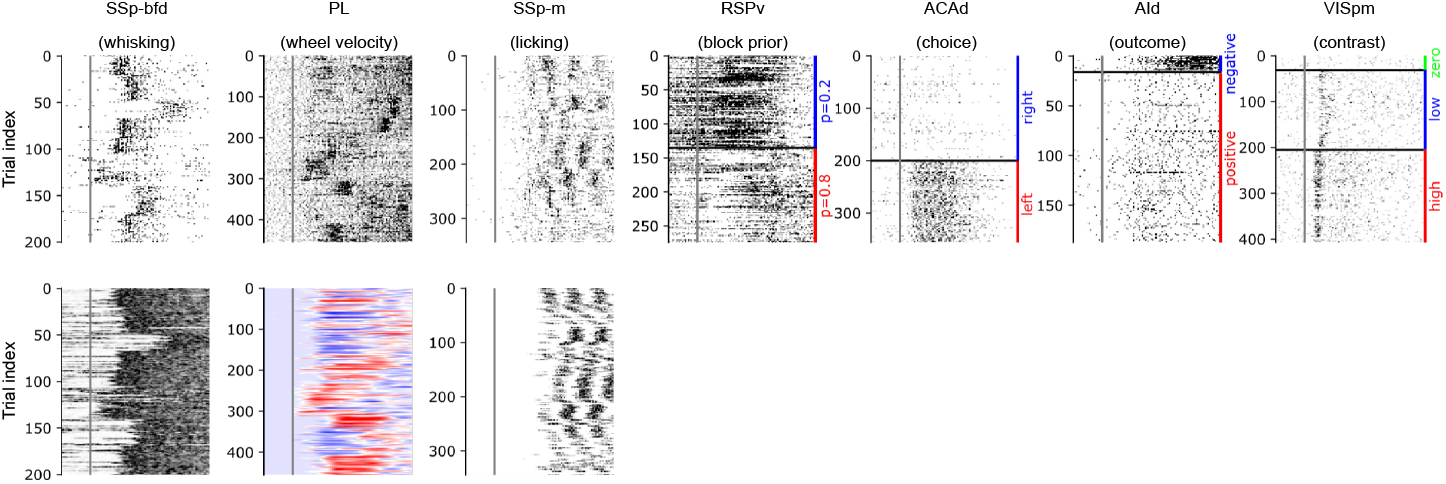
Neurons with high selectivity are strongly modulated by the corresponding input variable. Examples of neurons with strong selectivity to each input variable are displayed: the first row shows neuronal activity, while the second row illustrates the associated behavioral movements.

**Supplementary Figure 3.**
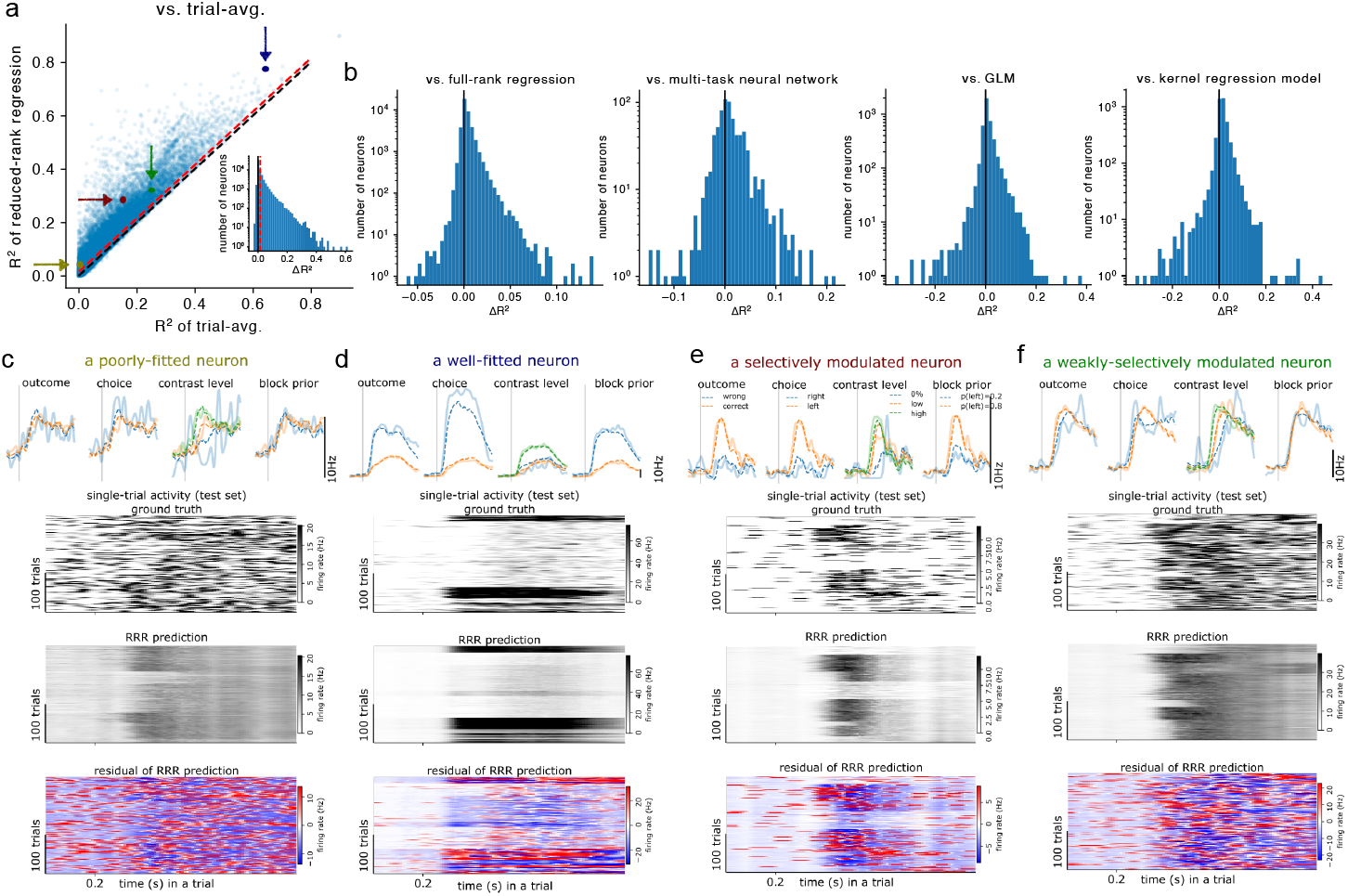
The reduced-rank regression (RRR) encoding model effectively reduces trial-to-trial variability and captures task-related variance. **a** Goodness-of-fit analysis comparing the RRR encoding model to the trial-averaged neural activity across all trials (i.e., the null model, Eq 8). Each point represents a neuron. The black line marks equal performance for both methods, while the red line indicates cases where RRR outperforms the trial-average estimate (threshold: 0.015). The four colored dots highlight the neurons shown in (c-f). **b** Comparison of RRR performance relative to other baseline models, measured as Δ*R*^2^ = *R*^2^(*RRR*)*−R*^2^(baseline method). Baseline models include full-rank regression (which assumes a linear encoding model without rank constraints), a multi-task neural network (a nonlinear model with feed-forward and recurrent layers as in [74]), a generalized linear model (GLM, adapted from [4]), and a kernel regression model (adapted from [21]). **c-f** Example neurons illustrating different response characteristics and how well RRR captures their activity. Top: Peri-stimulus time histograms (PSTHs) for different task conditions (color-coded). Middle: Single-trial activity (test set) alongside RRR predictions. Bottom: Residuals of the RRR predictions. **c** A neuron with low *R*^2^, where a large portion of neural variability is unexplained by RRR and not clearly linked to task variables. **d** A neuron with high *R*^2^, where RRR successfully captures most of the task-related variance. **e** A selectively modulated neuron, showing strong dependence on task conditions, reflected in both the PSTH and single-trial activity. RRR effectively models this neuron’s responses. **f** A weakly-selectively modulated neuron. Unlike (e), this neuron does not show strong differences in PSTHs across task conditions. However, a consistent step-like pattern appears in all PSTHs shortly after stimulus onset, suggesting that its response is primarily driven by task onset.

**Supplementary Figure 4.**
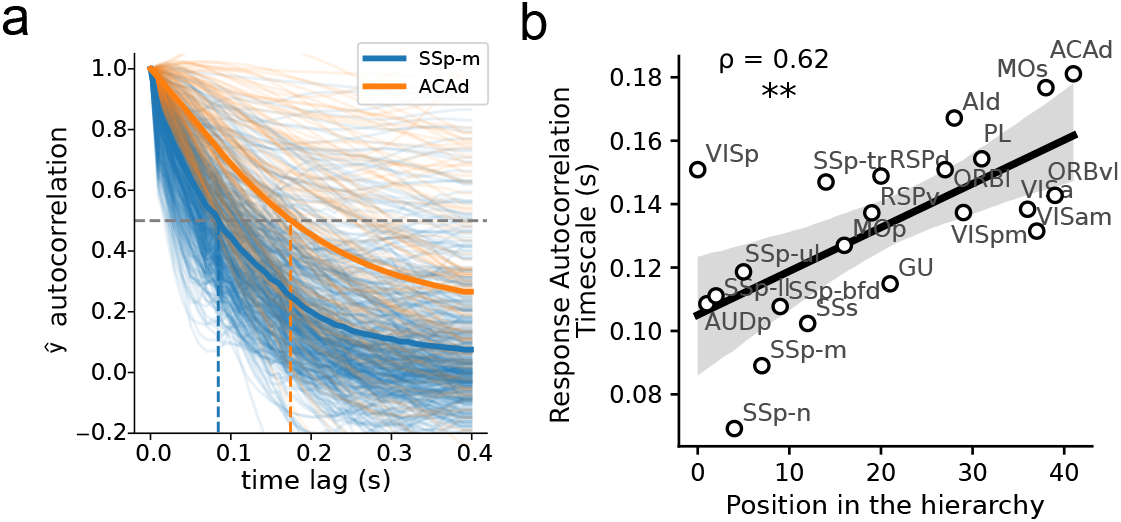
The analysis of responses autocorrelation times reveals a gradient along the cortical hierarchy. **a** The autocorrelation functions of single neurons’ responses to task variables estimated by the RRR (see Methods). Neurons from two example brain regions (SSp-m and ACAd) are shown, along with their mean profiles (solid orange and blue lines) and estimated autocorrelation timescales (orange and blue dashed lines). **b** Correlation between the position on the hierarchy of single cortical areas with their estimated autocorrelation timescales (see Methods).

**Supplementary Figure 5.**
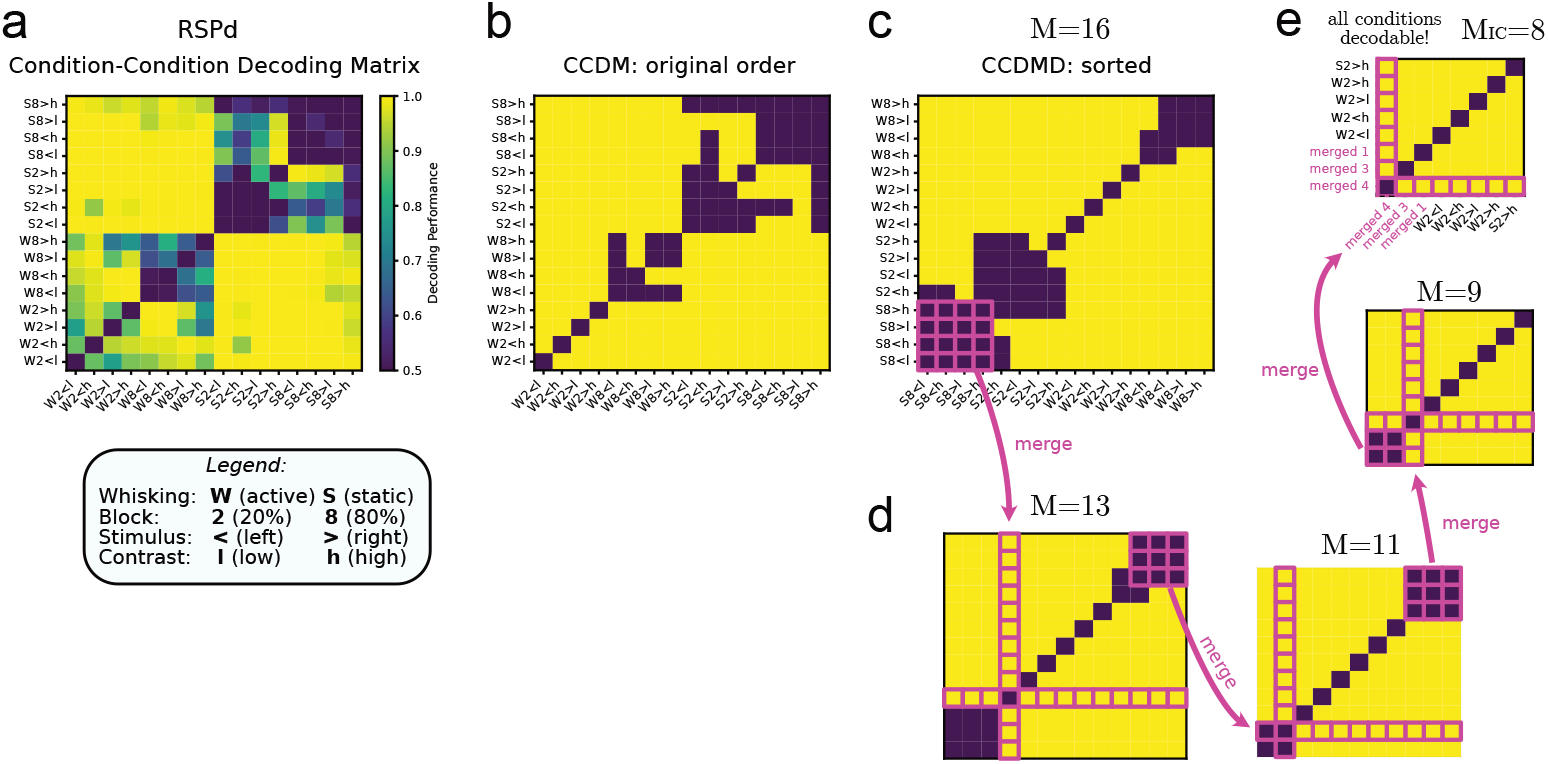
Example of all the individual steps of the independent conditions algorithm applied to the population activity of an example region (RSPd). First, a condition-condition decoding matrix (CCDM) is computed using cross-validated decoding with a linear SVC **(a)**, starting from the *M* = 16 conditions defined by the combinations of 4 binarized variables. The CCDM is then binarized into decodable and non-decodable pairs **(b)** by using a threshold (0.666). Then, the iterative algorithm finds the largest group of conditions that are non-decodable from each other, which corresponds to a dark block in the sorted matrix in **(c)**. These conditions are grouped together and given a new single-condition label **(d)**. The algorithm iterates until all the conditions are decodable from each other **(e)**. The size of the resulting CCDM is the number of independent conditions *M*_*IC*_.

**Supplementary Figure 6.**
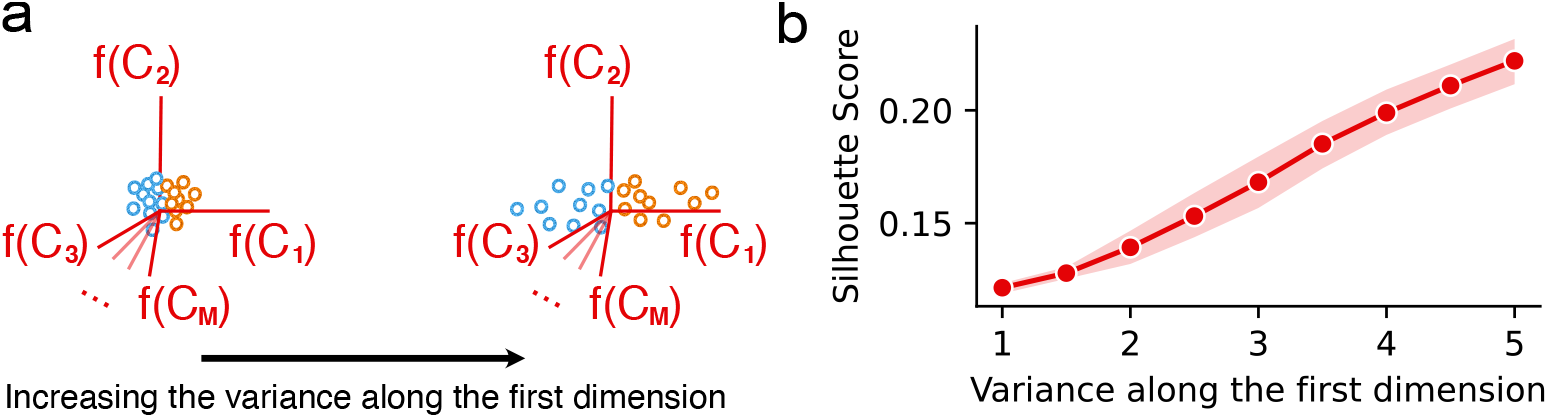
Effect of uneven data distribution on the silhouette score. (a) Schematic illustration of datasets with increasing variance along the first dimension, parameterized by *σ*_1_ in the covariance matrix *diag*(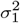, 1, *· · ·*, 1). Each dataset comprises 100 samples drawn from an eight-dimensional multivariate Gaussian distribution. (b) The silhouette score increases monotonically with *σ*_1_, showing that elongation of the data distribution can spuriously inflate the silhouette score even in the absence of genuine cluster structure. Shaded regions indicate the 95% confidence interval across independent realizations.

**Supplementary Figure 7.**
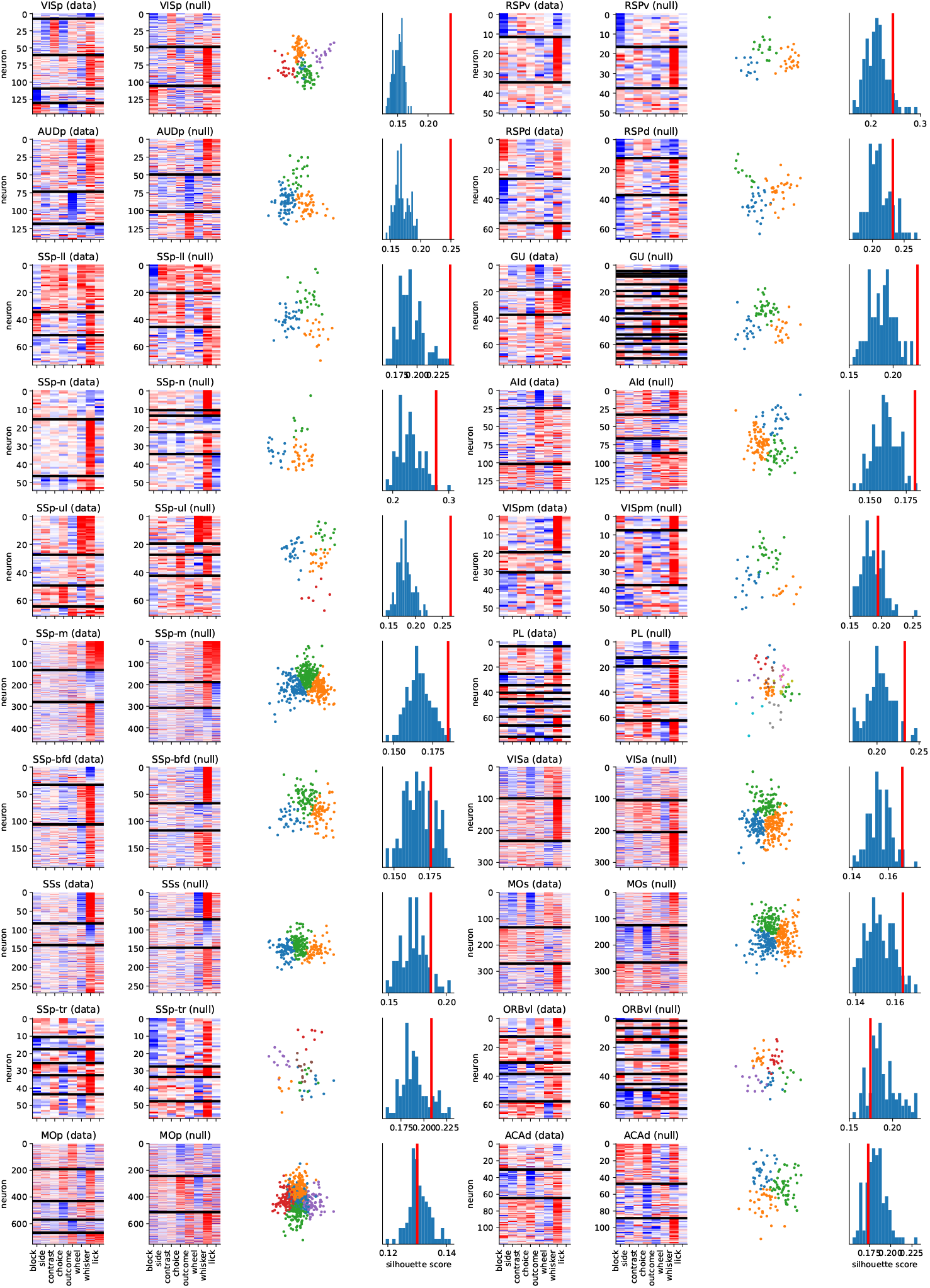
Clustering results of all cortical regions in the selectivity space. Subfigures are presented in the same format as Fig. 3bc.

**Supplementary Figure 8.**
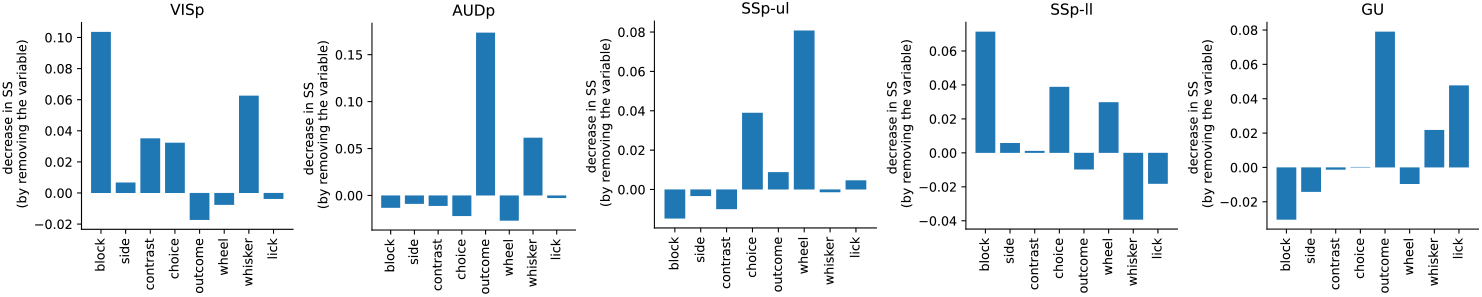
Key variables influencing clustering quality varied across brain areas and were often multi-modal, encompassing cognitive, movement, and sensory domains. To measure the contribution of individual variables to categorical clustering, we computed the difference in silhouette score found by k-means when removing each input variable. The five regions with the highest z-scored silhouette scores are shown.

**Supplementary Figure 9.**
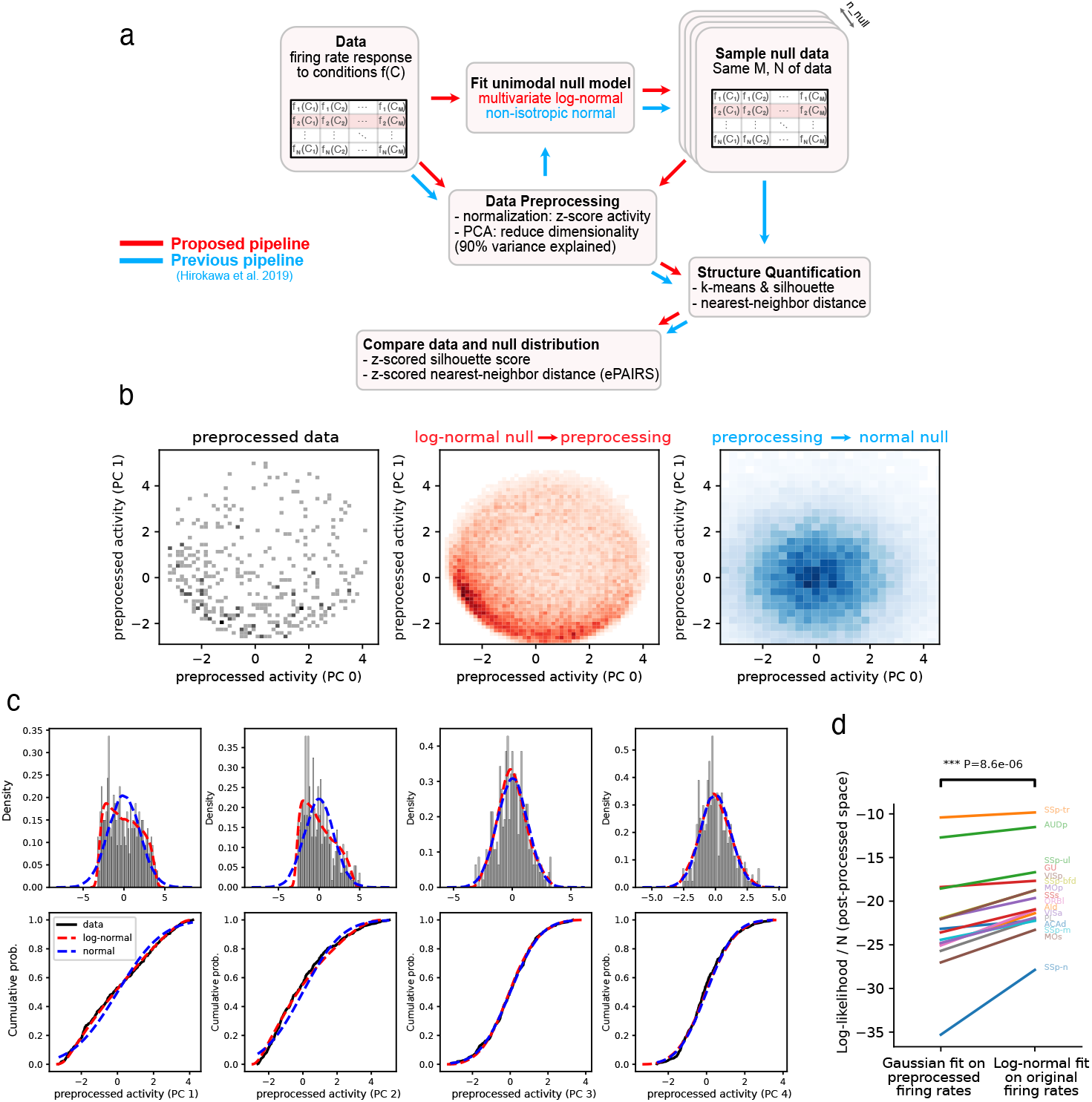
Pipeline comparison for analysis of structure in the conditions space. **(a)** Flow charts for the analysis of structure in the conditions space defined by the mean firing rates of neurons in response to a given set of experimental conditions, comparing the proposed pipeline with the pipeline developed in [2] for the ePAIRS null model comparison (blue arrows). The main methodological difference is that, in our pipeline, the uni-modal null model is fitted on the original data instead of the z-scored and dimensionality-reduced firing rate profiles. Note that in both cases, structural measures such as clustering quality or ePAIRS are computed in the post-processed, z-scored space. **(b)** Comparison of the joint distribution of the post-processed z-scored data (black) compared with the null model inferred using our pipeline (red: log-normal fitting in the original space, then z-scoring and dimensionality reduction) or using the pipeline as described in [2] (blue: z-scoring and reducing the data dimensionality and *then* fitting a Gaussian null model), for an example region (MOs). **(c)** Marginal distributions of the data along the first 4 principal components of the reduced-dimensionality z-scored conditions space (that in which we compute structure). Top row: distribution of the data (black) compared with the null model distribution given by our pipeline (red) and the pipeline in [2] (blue). Bottom row: cumulative distributions of the data in the top row (same color scheme). Note that, in this data, the distributions of the data along the first dimensions of the reduced-dimensionality space (noted as PC0 and PC1) are typically asymmetric and non-Gaussian. The asymmetry is well-captured by a log-normal null model in the original space that undergoes the same preprocessing of the data (red curve), while it is less well-captured by a post-processed Gaussian fit (blue curve). In this example, PCs after the second are more Gaussian. **(d)** Comparison of the goodness of fit, quantified as the normalized log-likelihood of the data given the fitted null distribution, between the two pipelines. p-value computed via a paired t-test.

**Supplementary Figure 10.**
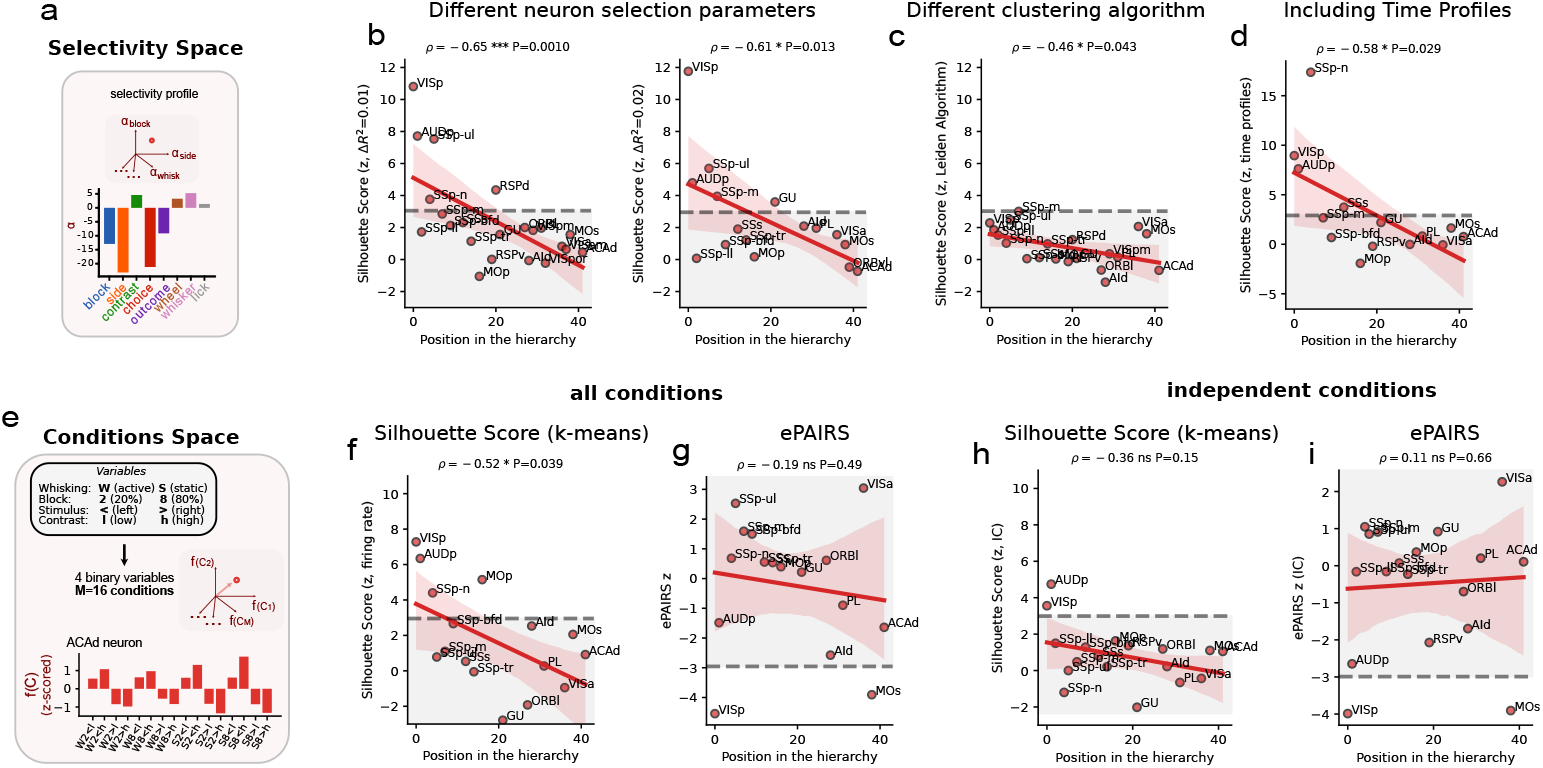
Additional analysis of structure in the selectivity and condition firing rate space. Comprehensive clustering analyses show consistent patterns in both selectivity space (a-d) and condition mean firing rate space (e-i) in agreement with the results in Fig. 3. **(a-d)** Analysis of structure in the selectivity space with different parameters and algorithms. **(a)** Visualization of an example neuronal selectivity profile in the selectivity space. (b-d) Same analysis as Fig. 3d (correlation of clustering quality in the selectivity space with the position of the hierarchy) reproduced with **(b)** different neuron selection Δ*R*^2^ thresholds, **(c)** a different algorithm than k-means for finding the clusters (Leiden clustering algorithm), and **(d)** expanding the selectivity space by including the time-varying selectivity profiles (see Methods). **(e-i)** Results of structure analysis in the conditions mean firing rate space using the analysis pipeline explained in Suppl. Fig. 9. **(e)** Illustration of the conditions space framework incorporating four binary variables resulting in 16 experimental conditions. **(f)** Analysis of clustering quality in the 16-dimensional mean firing rate space of the 16 experimental conditions defined by the combinations of the four selected variables (see panel **e** and main text). **(g)** Structure quantified by the ePAIRS analysis (mean nearest-neighbor distance compared to a null model through z-scoring, see Methods) in the same conditions mean firing rate space of panel **f. (h, i)** Same analyses of (f, g) in the space defined by the mean firing rate of independent conditions of each individual area.

**Supplementary Figure 11.**
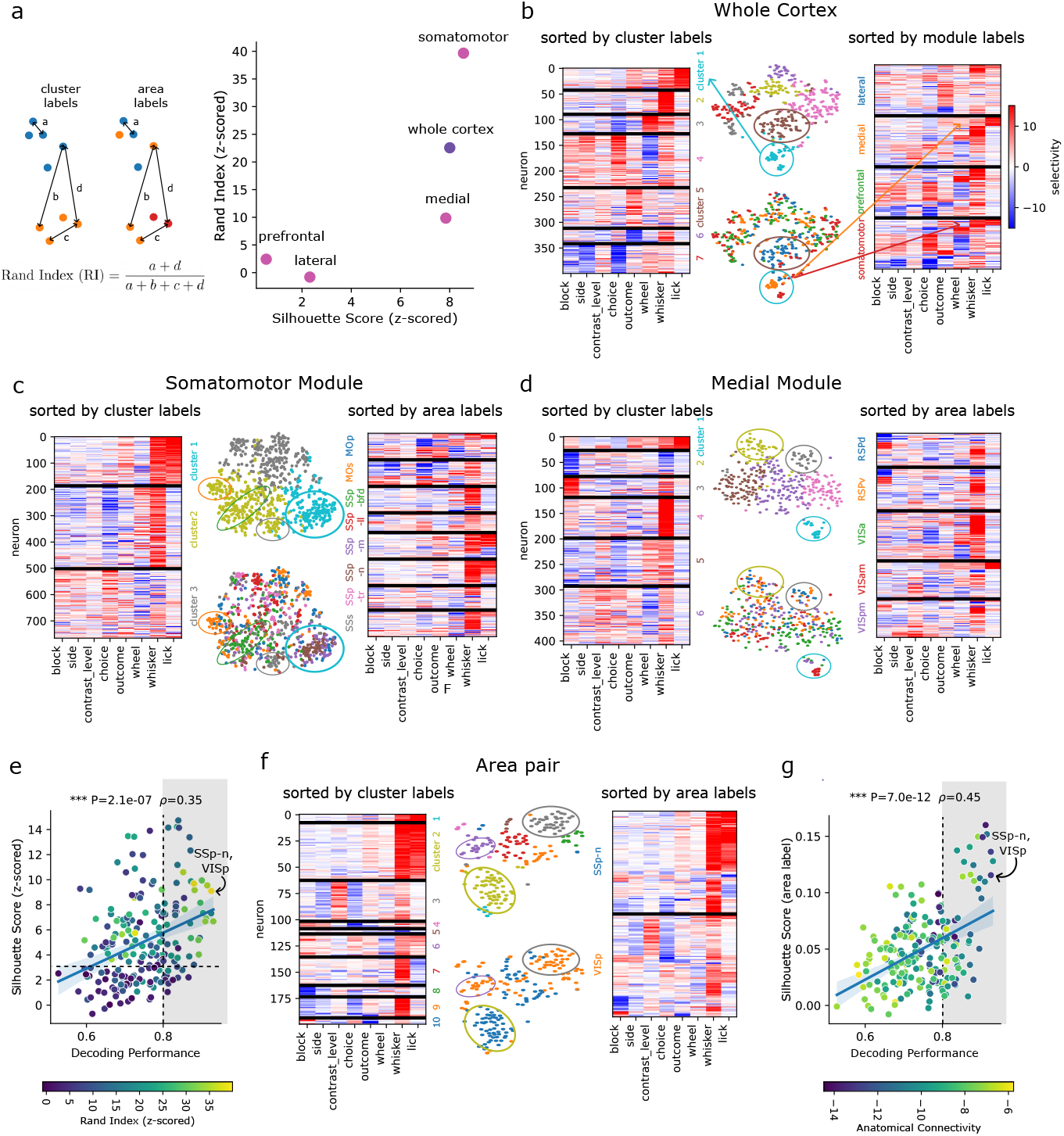
A globally organized brain, with functional clustering closely reflecting anatomical structure. **a** We propose a measure of how well neurons are clustered by anatomical structure using the rand index (z-scored). Left: The rand index (RI) quantifies the similarity between two clusterings. We compare the clustering based on neuron area labels to the optimal clustering solution. Right: Neurons in clustered modules (somatomotor and medial) and across the whole cortex are significantly organized by anatomical structure labels. **b** Neuron selectivity matrix with neurons pooled across the cortex, sorted either by cluster labels (left) or module labels (right), noted aside and distinguished with colors. Middle: t-SNE visualizations of the same matrix, colored by cluster (top) or module (bottom). As arrows illustrate, the full selectivity profile can be traced through the colors, while ellipses highlight examples of clustering by anatomical areas. Cluster 1, with strong selectivity for licking, for example, is primarily from somatomotor and medial modules. When analyzing a single population spanning the entire cortex, we observed categorical representations (*SS z ≃* 8), with key variables contributing to cluster formation encompassing nearly all variables considered. **c-d** Somatomotor (c) and medial (d) modules show clustering by areas. In the somatomotor module, clustering reflects differences in selectivity for whisking power and licking. Cluster 1 (high selectivity for both variables) is enriched in neurons from nose (SSp-n) and mouth (SSp-m) regions of primary somatosensory cortex. Cluster 2 (high whisking selectivity, low licking selectivity) contains neurons mainly from barrel field (SSp-bfd), secondary somatosensory cortex (SSs), and secondary motor cortex (MOs). Within this cluster, neurons from the same area are more similar to each other than to neurons from different areas. In the medial module, clustering is shaped by selectivity to block prior, whisking power, and licking. Cluster 1 (strong licking selectivity) is enriched in VISam, while clusters with strong block-prior selectivity are dominated by RSPd and RSPv neurons. **e** Area pairs with high decoding performance tend to show high z-scored SS and RI. The dashed horizontal line marks statistical significance (*p <* 0.05, Bonferroni corrected). The dashed vertical line and shaded area highlight pairs with high decoding performance (*>* 0.8). **f** Example area pair (VISp vs. SSp-n) showing clustering by areas. **g** Silhouette scores computed based on neuron area labels increase significantly with decoding performance. For clarity of the plots and to avoid biases toward areas or modules with larger neuron numbers, we considered the 100 best-encoded neurons from each module (b) or area (c, d, f).

**Supplementary Figure 12.**
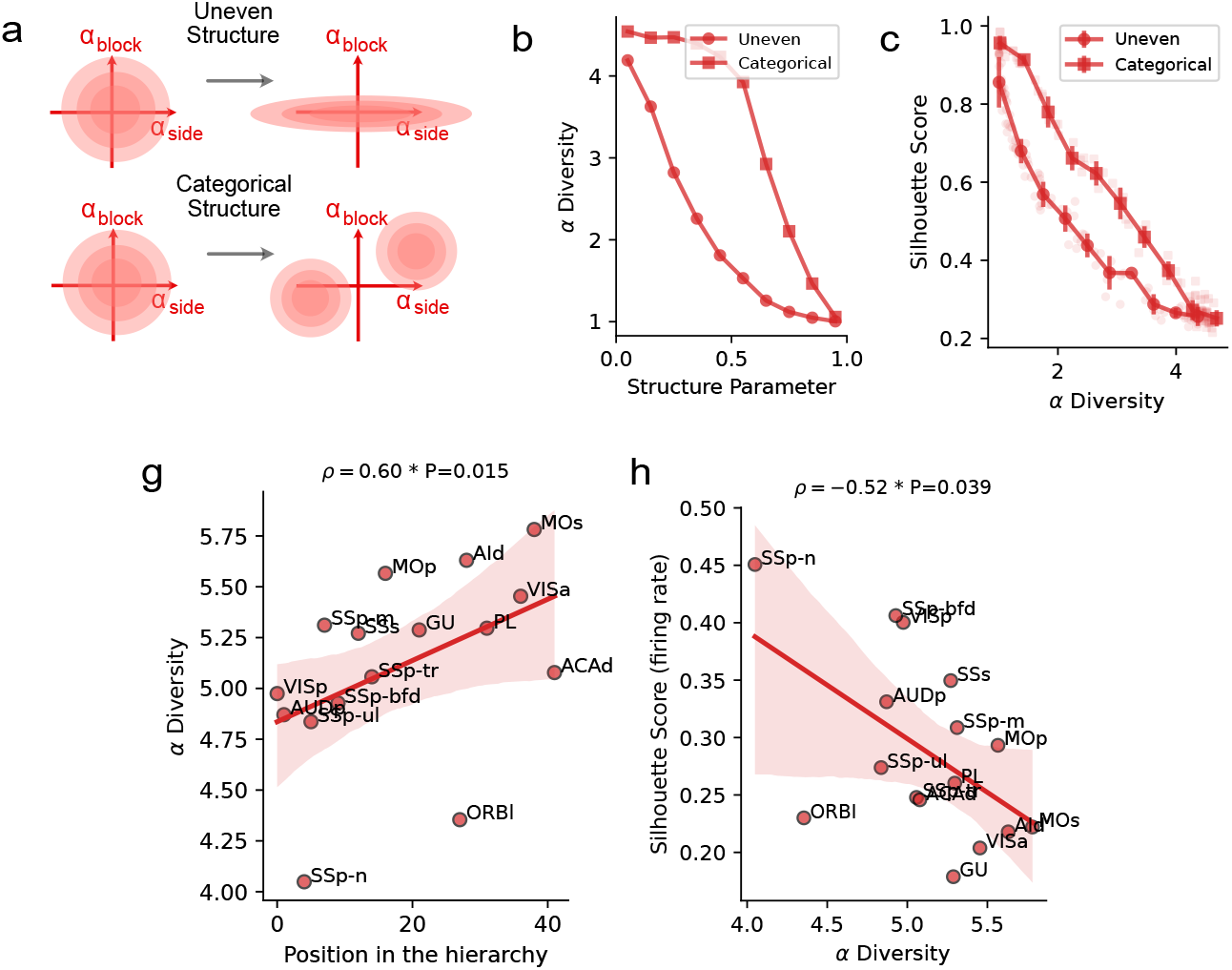
Synthetic exploration and additional analysis of *α*-diversity. **(a)** Conceptual model of how uneven and categorical structures shape selectivity profiles. In the simulations, structure is imposed in the space of regression coefficients *α* of neuronal response to a set of *V* binary variables, see Methods. Simulations are parametrized by a structure parameter that controls how strong these structures are. When this parameter is zero, alpha coefficients are maximally unstructured (random Gaussian). When the parameter is close to 1, alphas are maximally structured (categorical: perfect clusters; uneven: just one variable dominating the distribution). See Methods. **(b)** *α*-diversity decreases with structure in the alpha space, as expected. **(c)** Simulations showing how Silhouette Score changes with *α*-diversity for the two models. Points show individual simulations, while error bars show mean and standard deviation over a running window of 10% of the x-range. We used *γ* = 0.25, *k* = 2, *V* = 4, *N* = 100 (see Methods). **(g)** *α*-diversity increases along the cortical hierarchy. **(h)** *α*-diversity is inversely correlated to the silhouette score in the conditions firing rate space.

**Supplementary Figure 13.**
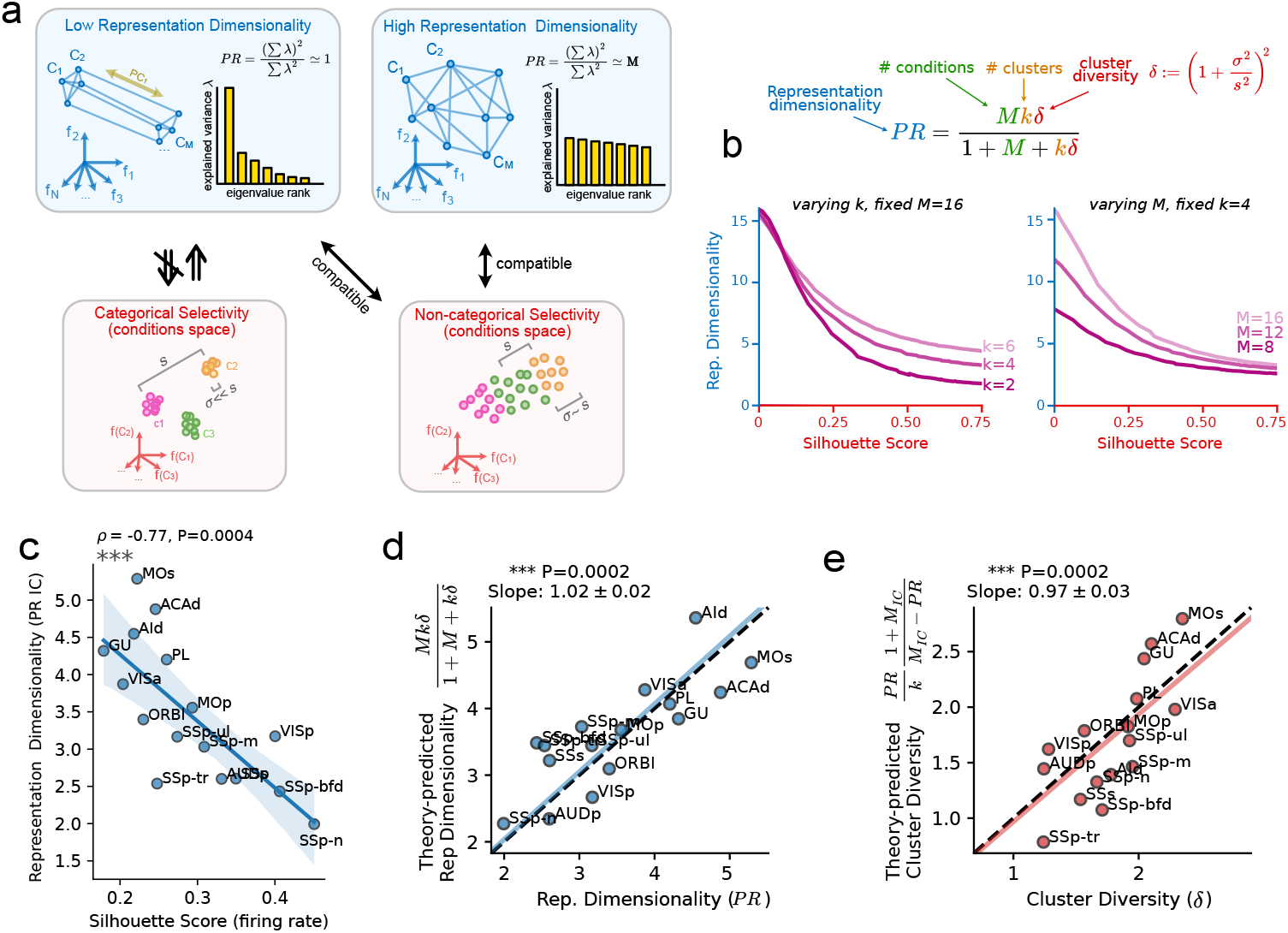
Theoretical Investigation of the relation between clustering of neural response profiles and embedding dimensionality of neural representations. **(a)** Schematic of how categorical versus non-categorical selectivity in the condition space shapes representational dimensionality. Categorical clusters in the conditions space imply a low-dimensional geometry, while non-categorical responses allow for a high-dimensional geometry. Note that low-dimensional geometries are compatible with non-categorical representations (see [3]), as long as the embedding dimensionality is the same across the two spaces. The intra-clustering diversity *δ* is quantified as the ratio of the dispersion of points within each cluster (denoted as *σ*) to the distance between different clusters (denoted as *s*). **(b)** Top: analytical relation between the features of a set of clusters in the conditions space (number of clusters *k*, number of conditions *M*, intra-cluster diversity *δ*) and their Representation dimensionality (PR) in the neural space. Bottom: visualization of how the Representation dimensionality (PR) is limited by clustering quality and how this limit varies with the number of clusters *k* (left panel) and the number of conditions *M* (right panel). For these visualizations, intra-clustering diversity *δ* was converted into a silhouette score to aid comparison with the data (see Methods). **(c)** Coherently with the theoretical model, Representation dimensionality decreases with increasing Silhouette Score in the conditions space (Spearman’s *ρ*=–0.77, P = 0.0004). **(d)** Correlation between the value for *PR* predicted by the theory, using *k, δ* and *M*_*IC*_ (y-axis) and that computed from the centroids of the *M*_*IC*_ conditions in the neural space. The blue line is the best fit zero-intercept slope, while the dashed black line is the identity function *y* = *x*. **(e)** Similar to **d**, the intra-cluster diversity *δ* in the conditions space could also be quantitatively predicted from *M*_*IC*_, *PR*, and *k* using an inversion of the equation in **b**.

**Supplementary Figure 14.**
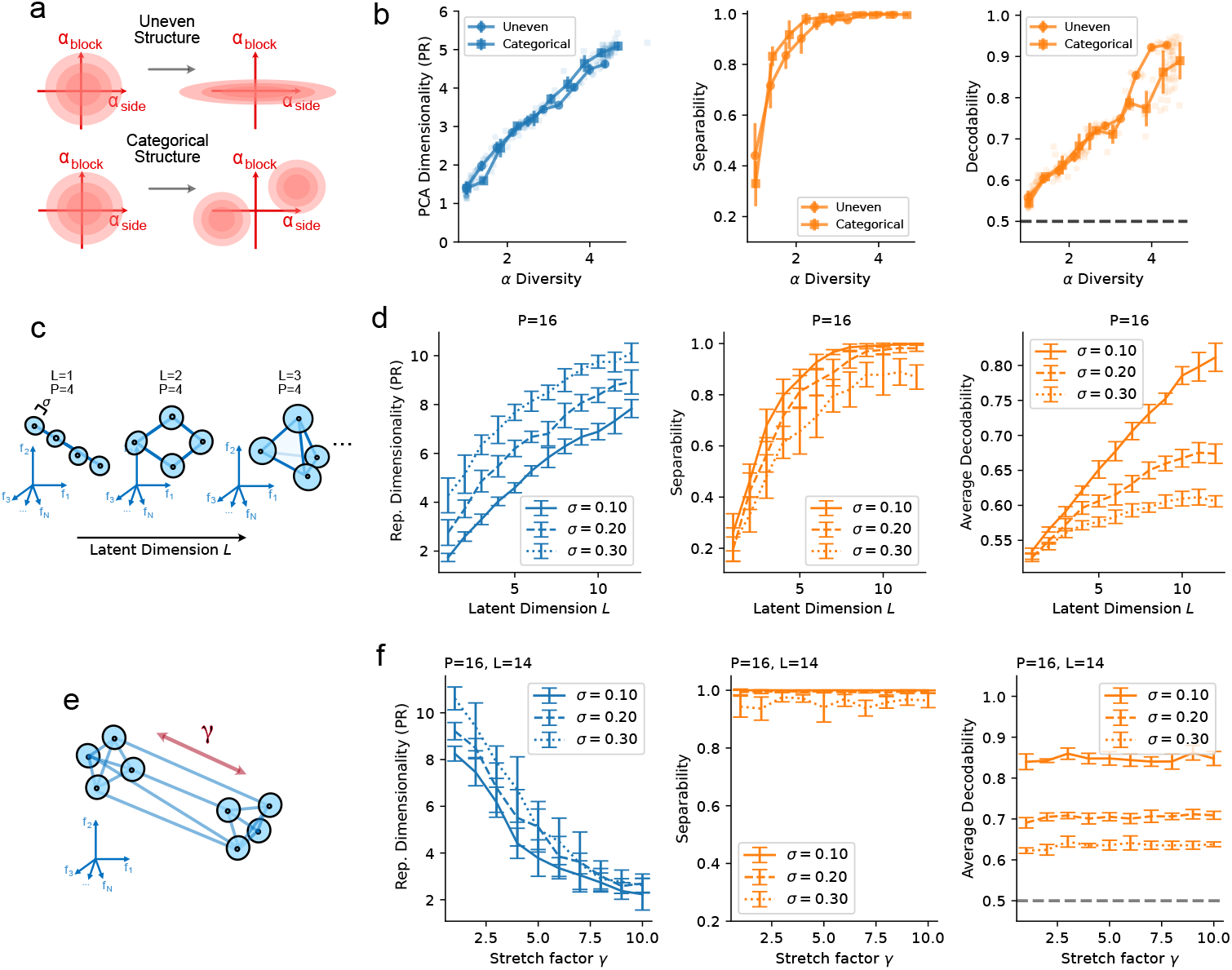
Synthetic exploration of how Representation dimensionality, Separability, and Average Decodability depend on diversity in the alpha space and dimensionality in the activity space. **(a)** Conceptual model of how uneven and categorical structures shape selectivity profiles. In the simulations, structure is imposed in the space of regression coefficients *α* of neuronal response to a set of *V* binary variables, see Methods. **(b)** Simulations showing how Representation Dimensionality, Separability, and Average Decodability change with *α*-diversity for the two models. Points show individual simulations, while error bars show mean and standard deviation over a running window of 10% of the x-range. We used *γ* = 0.25, *k* = 2, *V* = 4, *N* = 100 (see Methods). **(c)** Schematic of the subspace dimensionality analysis: in these simulations, P centroids are randomly sampled from an L-dimensional subspace of the N-dimensional activity space (see Methods). Trial-to-trial variability is then added to the centroids with a standard deviation *σ* (scaled with 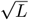 to keep signal-to-noise invariant with L) in the full N-dimensional space. **(d)** Representation Dimensionality, Separability, and Average Decodability as a function of L for different values of the trial-to-trial noise. Low-dimensional (*L* ≲ *M/*2 = 8) geometries limit the Separability, which saturates at larger dimensionalities. This behavior is observed at all different levels of noise. Decodability, on the other hand, increases almost linearly with L for small noise levels, and sub-linearly with medium and high noise. Each error bar shows mean and standard deviation across 10 simulations with fixed level of noise and latent dimensionality L. These simulations were performed using N=100, P=16, T=20. **(e)** While Representation dimensionality and Separability are genereally linked, this relationship is not 1-1. It is indeed possible to construct a geometry that has maximal separability *and* low Representation dimensionalities, by starting from a high-dimensional geometry and stretching one particular dimension while keeping the noise level unchanged. **(f)** This operation will decrease the Representation dimensionality while keeping separability intact. Decodability is also largely invariant upon this transformation (and is mostly determined by the noise level).

**Supplementary Figure 15.**
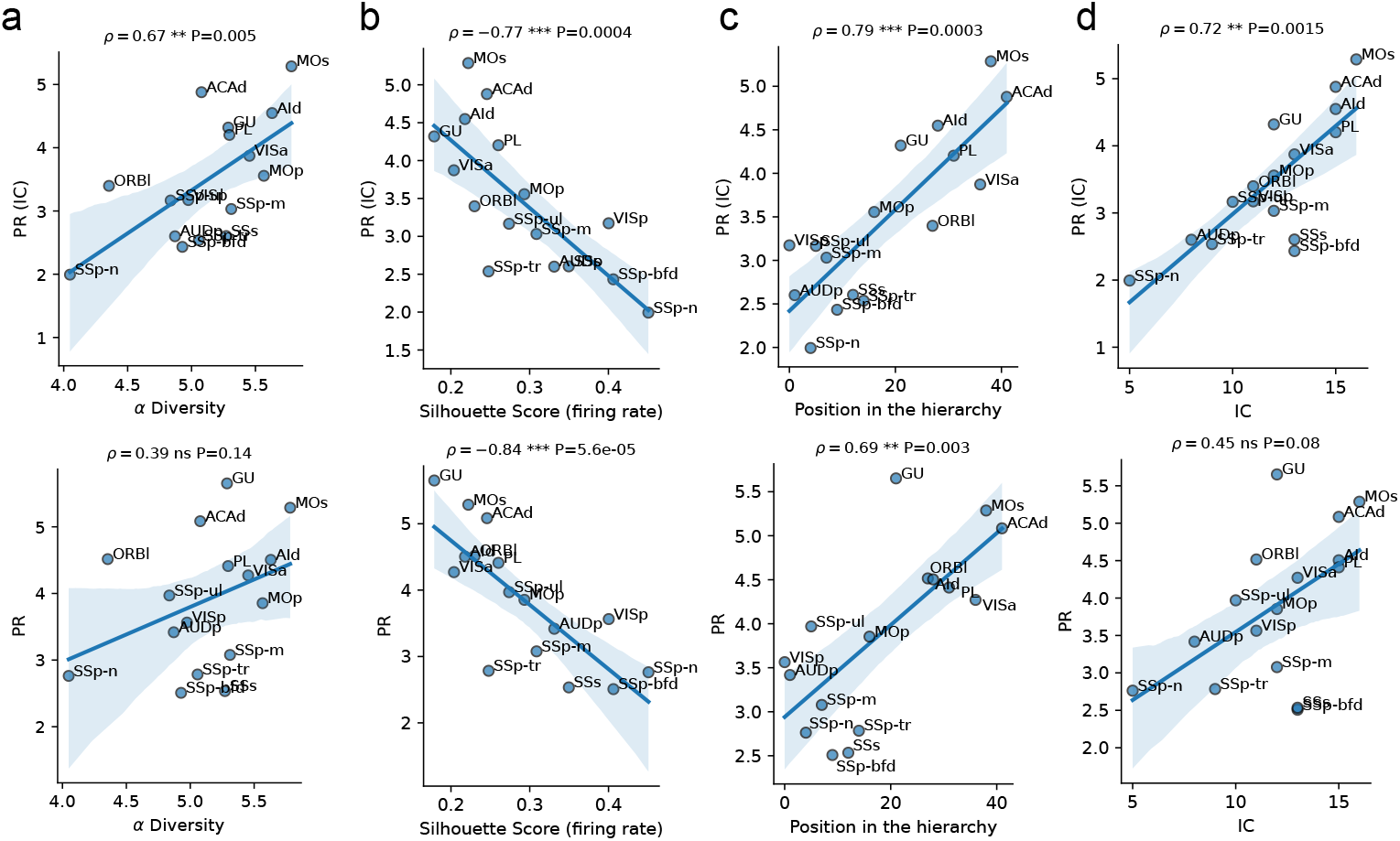
**a)** Representation dimensionality computed on independent conditions increases with *α−*diversity (top). PR of all conditions exhibit a non-significant trend in the same direction (bottom). **b)** Both Representation dimensionality computed on independent conditions (top, same data as Fig. 5c) and on all conditions (bottom) are inversely related to the Silhouette Score. **c)** Both measures increase along the cortical hierarchy (top: same data as Fig. 5d). **d)** Representation dimensionality of independent conditions increases with the number of independent conditions (top); a similar trend (non-significant) is observed for Representation dimensionality of all conditions.

**Supplementary Figure 16.**
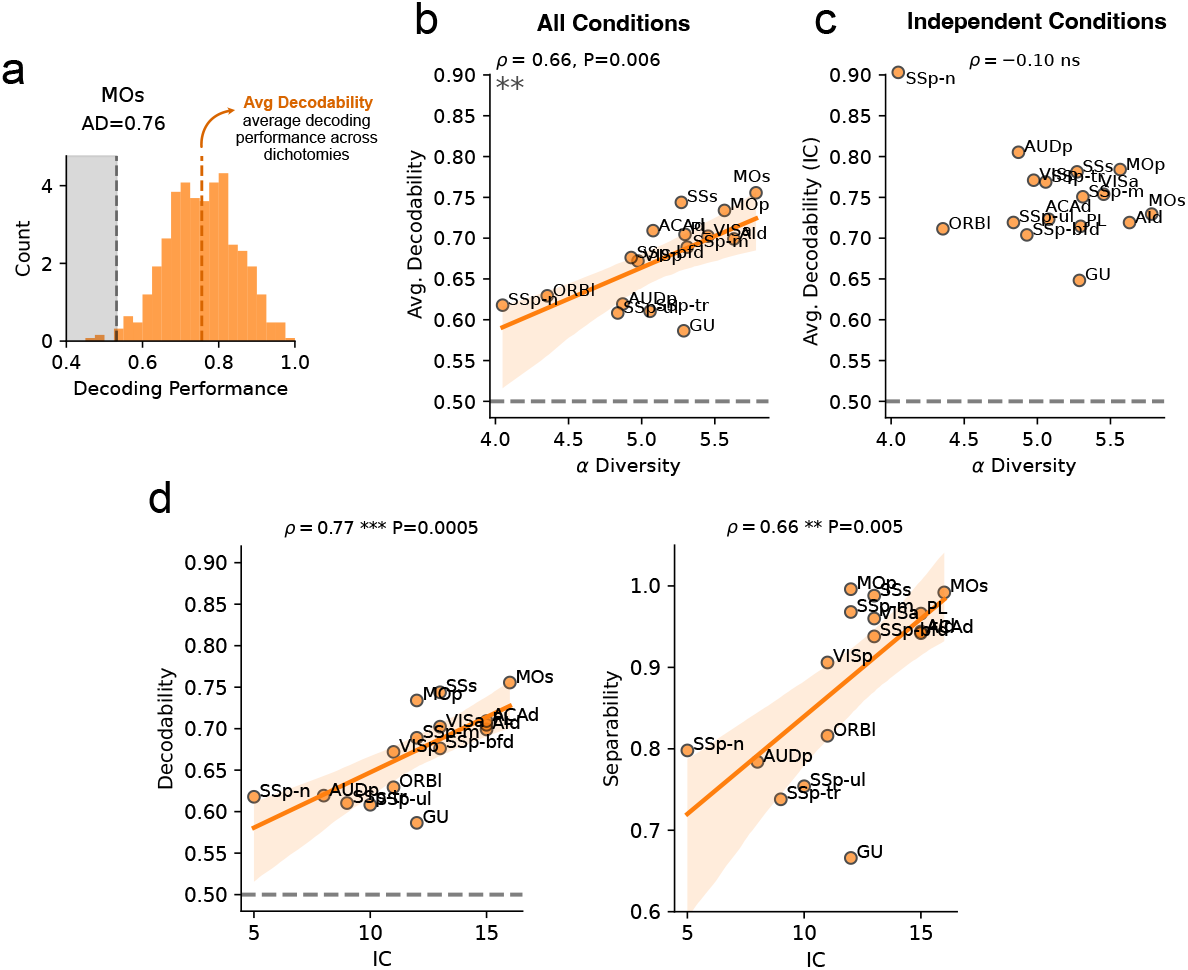
**(a)** Distribution of linear decoding performance across *n* = 200 random dichotomies of the *M* = 16 conditions in MOs (same data of Fig. 6b). Average Decodability (AD) is defined as the mean of this distribution. **(b)** AD increases with *α*-diversity when computed on all conditions. **(c)** This relationship is not present when computing AD for independent conditions only. **(d)** Across cortical areas, both AD and Separability are positively correlated with the number of independent conditions.

**Supplementary Figure 17.**
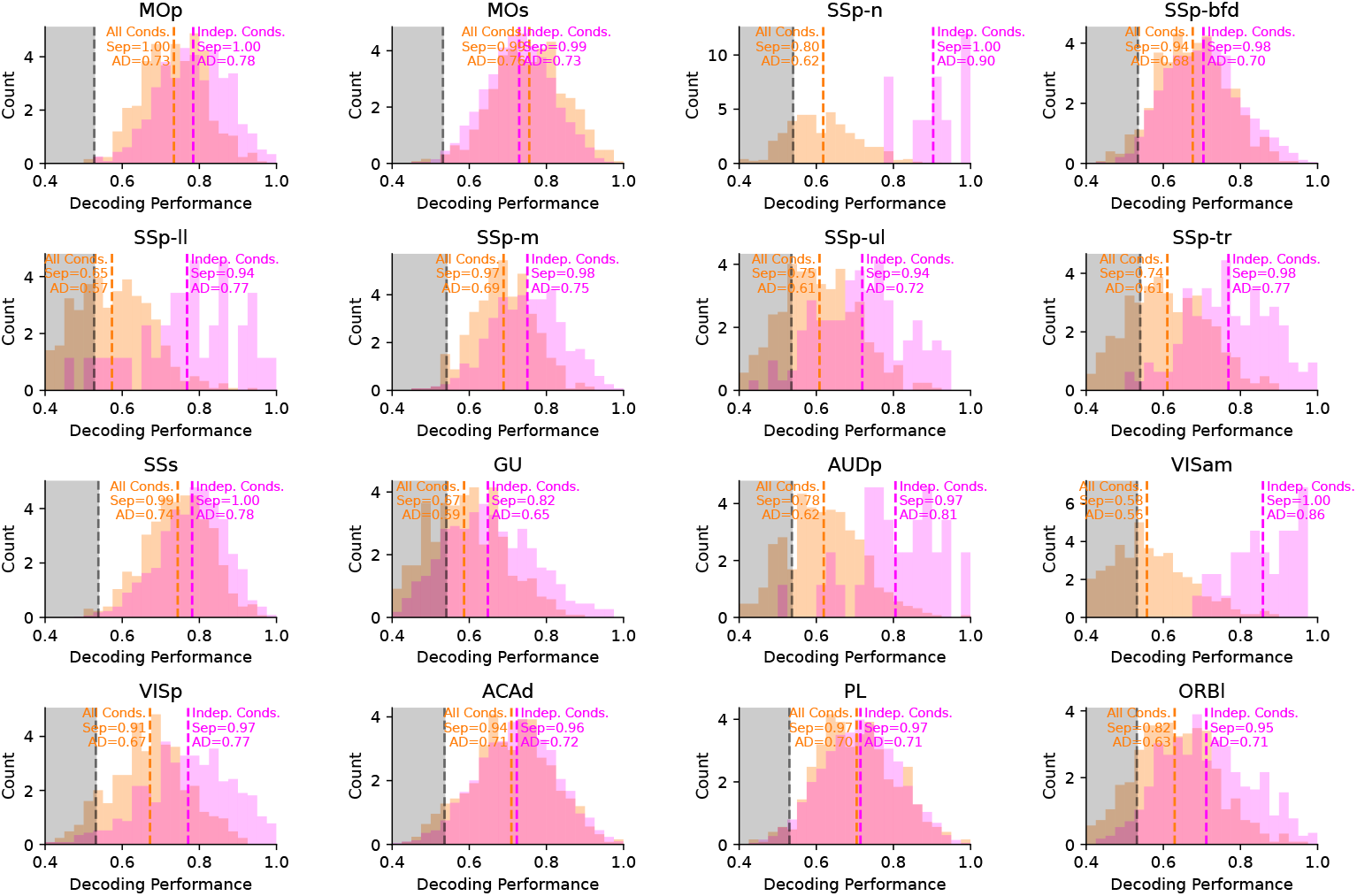
Distribution of decoding performance of *n* = 200 random dichotomies of all conditions (orange) and independent conditions (magenta) for all the regions used in Main Fig.s 5 and 6. Each performance was obtained by a cross-validated analysis of the decoding performance of a random balanced binary classification of the independent conditions found for each region. The cross-validation was performed on mean firing rate vectors corresponding to individual trials. Within each analysis, individual conditions were balanced within each side of the dichotomy to avoid grouped or large conditions dominating the analysis (see Methods). Separability was then computed as the fraction of dichotomy decoding performances that was larger than a null distribution found by shuffling the labels of individual trials, while Average Decodablility (AD) was computed as the mean decoding performance across the sampled random dichotomies.

## Footnotes

1 Here we considered neural responses over a different time window from [4] and enriched set of behavior movements.

2 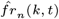 is calculated by inversing the z-score transformation applied in the preprocessing step 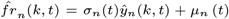 (Suppl. Fig. 1b-d).

## References

[1] Raposo, D., Kaufman, M. T. & Churchland, A. K. A category-free neural population supports evolving demands during decision-making. Nature neuroscience 17, 1784–1792 (2014).

[2] Hirokawa, J., Vaughan, A., Masset, P., Ott, T. & Kepecs, A. Frontal cortex neuron types categorically encode single decision variables. Nature 576, 446–451 (2019).

[3] Kaufman, M. T. et al. The implications of categorical and category-free mixed selectivity on representational geometries. Current opinion in neurobiology 77, 102644 (2022).

[4] Angelaki, D. et al. A brain-wide map of neural activity during complex behaviour. Nature 645, 177–191 (2025).

[5] Harris, J. A. et al. Hierarchical organization of cortical and thalamic connectivity. Nature 575, 195–202 (2019).

[6] Rigotti, M. et al. The importance of mixed selectivity in complex cognitive tasks. Nature 497, 585–590 (2013).

[7] Fusi, S., Miller, E. K. & Rigotti, M. Why neurons mix: high dimensionality for higher cognition. Current opinion in neurobiology 37, 66–74 (2016).

[8] Tye, K. M. et al. Mixed selectivity: Cellular computations for complexity. Neuron (2024).

[9] Ostojic, S. & Fusi, S. Computational role of structure in neural activity and connectivity. Trends in Cognitive Sciences (2024).

[10] Hocker, D. L., Brody, C. D., Savin, C. & Constantinople, C. M. Subpopulations of neurons in lofc encode previous and current rewards at time of choice. Elife 10, e70129 (2021).

[11] Onken, A., Xie, J., Panzeri, S. & Padoa-Schioppa, C. Categorical encoding of decision variables in orbitofrontal cortex. PLoS computational biology 15, e1006667 (2019).

[12] Blanchard, T. C., Piantadosi, S. T. & Hayden, B. Y. Robust mixture modeling reveals category-free selectivity in reward region neuronal ensembles. Journal of neurophysiology 119, 1305–1318 (2018).

[13] Hardcastle, K., Maheswaranathan, N., Ganguli, S. & Giocomo, L. M. A multiplexed, heterogeneous, and adaptive code for navigation in medial entorhinal cortex. Neuron 94, 375–387 (2017).

[14] Bernardi, S. et al. The geometry of abstraction in the hippocampus and prefrontal cortex. Cell 183, 954–967 (2020).

[15] Boyle, L. M., Posani, L., Irfan, S., Siegelbaum, S. A. & Fusi, S. Tuned geometries of hippocampal representations meet the computational demands of social memory. Neuron 112, 1358–1371 (2024).

[16] O’Neill, P.-K. et al. The representational geometry of emotional states in basolateral amygdala. bioRxiv 2023–09 (2023).

[17] Courellis, H. S. et al. Abstract representations emerge in human hippocampal neurons during inference. Nature 1–9 (2024).

[18] Jazayeri, M. & Ostojic, S. Interpreting neural computations by examining intrinsic and embedding dimensionality of neural activity. Current opinion in neurobiology 70, 113–120 (2021).

[19] Khosla, M., Williams, A. H., McDermott, J. & Kanwisher, N. Privileged representational axes in biological and artificial neural networks. bioRxiv 2024–06 (2024).

[20] Pillow, J. W. et al. Spatio-temporal correlations and visual signalling in a complete neuronal population. Nature 454, 995–999 (2008).

[21] Steinmetz, N. A., Zatka-Haas, P., Carandini, M. & Harris, K. D. Distributed coding of choice, action and engagement across the mouse brain. Nature 576, 266–273 (2019).

[22] Murray, J. D. et al. A hierarchy of intrinsic timescales across primate cortex. Nature neuroscience 17, 1661–1663 (2014).

[23] Zeraati, R., Shi, Y., Levina, A., Engel, T. et al. A census of neural timescales across the mouse brain (2024).

[24] Song, M. et al. Hierarchical gradients of multiple timescales in the mammalian forebrain. bioRxiv 2023–05 (2023).

[25] Rudelt, L. et al. Signatures of hierarchical temporal processing in the mouse visual system. PLOS Computational Biology 20, e1012355 (2024).

[26] Gao, P. & Ganguli, S. On simplicity and complexity in the brave new world of large-scale neuroscience. Current opinion in neurobiology 32, 148–155 (2015).

[27] Rousseeuw, P. J. Silhouettes: a graphical aid to the interpretation and validation of cluster analysis. Journal of computational and applied mathematics 20, 53–65 (1987).

[28] Traag, V. A., Waltman, L. & Van Eck, N. J. From louvain to leiden: guaranteeing well-connected communities. Scientific reports 9, 1–12 (2019).

[29] Litwin-Kumar, A., Harris, K. D., Axel, R., Sompolinsky, H. & Abbott, L. Optimal degrees of synaptic connectivity. Neuron 93, 1153–1164 (2017).

[30] Dahmen, D. et al. Strong coupling and local control of dimensionality across brain areas. Biorxiv 2020–11 (2020).

[31] Rigotti, M., Rubin, D. B. D.Wang, X.-J. & Fusi, S. Internal representation of task rules by recurrent dynamics: the importance of the diversity of neural responses. Frontiers in computational neuroscience 4, 24 (2010).

[32] Cover, T. M. Geometrical and statistical properties of systems of linear inequalities with applications in pattern recognition. IEEE transactions on electronic computers 326–334 (2006).

[33] Boser, B. E., Guyon, I. M. & Vapnik, V. N. A training algorithm for optimal margin classifiers, 144–152 (1992).

[34] Dyballa, L. et al. Population encoding of stimulus features along the visual hierarchy. Proceedings of the National Academy of Sciences 121, e2317773121 (2024).

[35] Freedman, D. J. & Assad, J. A. Experience-dependent representation of visual categories in parietal cortex. Nature 443, 85–88 (2006).

[36] Sarma, A., Masse, N. Y.Wang, X.-J. & Freedman, D. J. Task-specific versus generalized mnemonic representations in parietal and prefrontal cortices. Nature neuroscience 19, 143–149 (2016).

[37] Diedrichsen, J. & Kriegeskorte, N. Representational models: A common framework for understanding encoding, pattern-component, and representational-similarity analysis. PLoS computational biology 13, e1005508 (2017).

[38] Zhang, M. et al. Molecularly defined and spatially resolved cell atlas of the whole mouse brain. Nature 624, 343–354 (2023).

[39] Markram, H. et al. Interneurons of the neocortical inhibitory system. Nature reviews neuroscience 5, 793–807 (2004).

[40] Bugeon, S. et al. A transcriptomic axis predicts state modulation of cortical interneurons. Nature 607, 330–338 (2022).

[41] Yu, H. et al. In vivo cell-type and brain region classification via multimodal contrastive learning. bioRxiv 2024–11 (2024).

[42] Stringer, C., Pachitariu, M., Steinmetz, N., Carandini, M. & Harris, K. D. High-dimensional geometry of population responses in visual cortex. Nature 571, 361–365 (2019).

[43] Recanatesi, S., Bradde, S., Balasubramanian, V., Steinmetz, N. A. & Shea-Brown, E. A scale-dependent measure of system dimensionality. Patterns 3 (2022).

[44] Hopfield, J. J. Neural networks and physical systems with emergent collective computational abilities. Proceedings of the national academy of sciences 79, 2554–2558 (1982).

[45] Amit, D. J. Modeling brain function: The world of attractor neural networks (Cambridge university press, 1989).

[46] Jaeger, H. & Haas, H. Harnessing nonlinearity: Predicting chaotic systems and saving energy in wireless communication. science 304, 78–80 (2004).

[47] Maass, W., Natschläger, T. & Markram, H. Real-time computing without stable states: A new framework for neural computation based on perturbations. Neural computation 14, 2531–2560 (2002).

[48] Buonomano, D. V. & Maass, W. State-dependent computations: spatiotemporal processing in cortical networks. Nature Reviews Neuroscience 10, 113–125 (2009).

[49] Barak, O., Rigotti, M. & Fusi, S. The sparseness of mixed selectivity neurons controls the generalization– discrimination trade-off. Journal of Neuroscience 33, 3844–3856 (2013).

[50] Babadi, B. & Sompolinsky, H. Sparseness and expansion in sensory representations. Neuron 83, 1213– 1226 (2014).

[51] Dominé, C. C. et al. From lazy to rich: Exact learning dynamics in deep linear networks. arXiv preprint 2409.14623 (2024).

[52] Tishby, N., Pereira, F. C. & Bialek, W. The information bottleneck method. arXiv preprint physics/0004057 (2000).

[53] Papyan, V., Han, X. & Donoho, D. L. Prevalence of neural collapse during the terminal phase of deep learning training. Proceedings of the National Academy of Sciences 117, 24652–24663 (2020).

[54] Recanatesi, S. et al. Dimensionality compression and expansion in deep neural networks. arXiv preprint 1906.00443 (2019).

[55] Long, B.Yu, C.-P. & Konkle, T. Mid-level visual features underlie the high-level categorical organization of the ventral stream. Proceedings of the National Academy of Sciences 115, E9015–E9024 (2018).

[56] Doshi, F. R. & Konkle, T. Cortical topographic motifs emerge in a self-organized map of object space. Science Advances 9, eade8187 (2023).

[57] Prince, J. S., Alvarez, G. A. & Konkle, T. Contrastive learning explains the emergence and function of visual category-selective regions. Science Advances 10, eadl1776 (2024).

[58] Chang, L. & Tsao, D. Y. The code for facial identity in the primate brain. Cell 169, 1013–1028 (2017).

[59] Muscinelli, S. P., Wagner, M. J. & Litwin-Kumar, A. Optimal routing to cerebellum-like structures. Nature neuroscience 26, 1630–1641 (2023).

[60] Whittington, J. C., Dorrell, W., Ganguli, S. & Behrens, T. E. Disentanglement with biological constraints: A theory of functional cell types. arXiv preprint 2210.01768 (2022).

[61] Yang, G. R., Joglekar, M. R., Song, H. F., Newsome, W. T. & Wang, X.-J. Task representations in neural networks trained to perform many cognitive tasks. Nature neuroscience 22, 297–306 (2019).

[62] Driscoll, L. N., Shenoy, K. & Sussillo, D. Flexible multitask computation in recurrent networks utilizes shared dynamical motifs. Nature Neuroscience 27, 1349–1363 (2024).

[63] Dubreuil, A., Valente, A., Beiran, M., Mastrogiuseppe, F. & Ostojic, S. The role of population structure in computations through neural dynamics. Nature neuroscience 25, 783–794 (2022).

[64] Johnston, W. J. & Fusi, S. Modular representations emerge in neural networks trained to perform context-dependent tasks. bioRxiv 2024–09 (2024).

[65] Raichle, M. E. & Gusnard, D. A. Appraising the brain’s energy budget. Proceedings of the National Academy of Sciences 99, 10237–10239 (2002).

[66] Chen, Y. T. & Witten, D. M. Selective inference for k-means clustering. Journal of Machine Learning Research 24, 1–41 (2023).

[67] Merre, P. L. et al. A prefrontal cortex map based on single neuron activity. bioRxiv 2024–11 (2024).

[68] Williams, A. H. et al. Discovering precise temporal patterns in large-scale neural recordings through robust and interpretable time warping. Neuron 105, 246–259 (2020).

[69] Izenman, A. J. Reduced-rank regression for the multivariate linear model. Journal of multivariate analysis 5, 248–264 (1975).

[70] Posani, L. Decodanda: a python package for decoding and geometrical analysis of neural activity. In preparation. Available on Github: https://www.github.com/lposani/decodanda (2024).

[71] Pedregosa, F. et al. Scikit-learn: Machine learning in python. the Journal of machine Learning research 12, 2825–2830 (2011).

[72] Bron, C. & Kerbosch, J. Algorithm 457: finding all cliques of an undirected graph. Communications of the ACM 16, 575–577 (1973).

[73] Langdon, C. & Engel, T. A. Embedding dimension of neural manifolds and the structure of mixed selectivity. In preparation. (2024).

[74] Laboratory, I. B. et al. Reproducibility of in-vivo electrophysiological measurements in mice. bioRxiv 2022–05 (2022).

